# Principles for Systematic Optimization of an Orthogonal Translation System with Enhanced Biological Tolerance

**DOI:** 10.1101/2021.05.20.444985

**Authors:** Kyle Mohler, Jack Moen, Svetlana Rogulina, Jesse Rinehart

## Abstract

Over the past twenty years, the development of orthogonal biological systems has sparked a revolution in our ability to study cellular physiology. Orthogonal translation systems (OTSs) enable site-specific incorporation of hundreds of non-standard amino acids, offering unprecedented access to the study of cellular mechanisms modulated by post-translational modifications (e.g. protein phosphorylation). Although development of phosphoserine-OTSs (pSerOTS) has been significant, little work has focused on the biology of OTS development and utilization. To better understand the impact of OTSs on host physiology, we utilize pSerOTS as a model to systematically explore the extent to which OTS components interact with *Escherichia coli*. Using this information, we constructed pSerOTS variants designed to enhance OTS orthogonality by minimizing interactions with host processes and decreasing stress response activation. Our expanded understanding of OTS:host interactions enables informed OTS design practices which minimize the negative impact of OTSs while improving OTS performance across a range of experimental settings.

## Introduction

Cellular processes have evolved to produce and maintain a finely tuned balance of macromolecular substrates which drive cellular function. These substrate pools are highly dynamic; responding quickly to environmental stimuli to mitigate stress and adapt to nutrient availability^1^. Many adaptation mechanisms are ultimately mediated by transcriptional reprogramming, but the first sensor of cellular stress is often revolves around a substrate pool, e.g. nucleotides or amino acids^2^. Due to the extreme metabolic cost of amino acid catabolism, the amino acid pool is one of the primary sensors used to monitor and report stress associated with nutrient depletion^3^. The level of an individual amino acid can directly impact the constituency of the amino acid pool through allosteric regulation and feedback inhibition of metabolic enzymes^1^. More generally, amino acid levels are monitored indirectly as substrates in the synthesis of aminoacyl-tRNAs^4^. Aminoacyl-tRNAs (aa-tRNAs) are produced by aminoacyl-tRNA synthetases (aaRSs) in a two-step reaction in which an amino acid is activated using ATP to form an aminoacyl-adenylate. The activated amino acid is then transferred to its cognate tRNA to form an aa-tRNA^5,6^.

As direct substrates for translation, aa-tRNAs provide an efficient method to detect any stress to the cell that negatively impacts protein translation. The bacterial stringent response, for example, integrates cues from several nutrient sensing mechanisms to mount a complex global response to nutritional stress by closely monitoring aa-tRNA pool aminoacylation status^2^. However, this leaves the cell susceptible to conditions which artificially perturb the balance of translational substrates pools. Disruption of amino acid pool equilibrium, specifically, can perturb the aa-tRNA pool, resulting in a decrease in translational fidelity by altering kinetic parameters of tRNA aminoacylation and typically leads to activation of the bacterial stringent response^7^. Similarly, changes in the activity or fidelity of aaRSs have been demonstrated to alter the ability of the cell to accurately sense amino acid starvation in *E. coli* and *S. cerevisiae^8–10^*.

The field of synthetic biology has fundamentally expanded the repertoire of translational machinery^11^. Advances in gene mining and directed evolution techniques have generated a wide range of aaRS variants enabling the site-specific incorporation of hundreds of non-standard amino acids (nsAAs)^12,13^. These orthogonal translation systems (OTSs) have been leveraged to enhance the functionality of enzymes via systematic incorporation of functionalized amino acid analogues to increase industrial value^14,15^. OTSs also provide unprecedented access to the study of biological systems modulated by post-translational modifications (PTMs). Many PTMs are reversible and often transient in nature making them difficult to study in their native contexts. Protein phosphorylation is one of the most heavily utilized PTMs in the cell, yet it’s dynamic regulation makes it extremely difficult to study^16,17^. Recent advances in the construction of OTSs which enable the site specific incorporation of phosphorylated amino acids, e.g. phosphoserine (pSer)^18^, phosphothreonine (pThr)^19^, and phosphotyrosine (pTyr)^20–22^, have provided unprecedented access to the study of protein phosphorylation. The most well validated phospho-OTS is the phosphoserine-OTS (pSerOTS)^23–26^. The pSerOTS uses a phosphoseryl-tRNA synthetase (pSerRS) derived from *Methanococcus meriplaudis* to aminoacylate pSer onto a tRNA^Cys^ from *Methanococcus janaschii* modified to create a UAG-decoding suppressor tRNA^pSer^ with specificity for pSerRS. To facilitate delivery of pSer-tRNA^pSer^ to the ribosome, *E. coli* elongation factor Tu (EF-Tu) was evolved to bind the larger, negatively charged moiety creating EF-pSer (**Figure 1A**)^18^. Since its initial deployment, several additional pSerOTS variants have been released with functional improvements aimed at enhancing purity and yield of modified recombinant protein production. Although the study of the pSerOTS itself has been significant, relatively little work focused on the host aspect of OTS deployment. A major advancement in the deployment of OTSs, in general, was the development of a recoded strain of *E. coli* (C321.ΔA)^27^. This strain was modified using multiplexed automatable genome engineering (MAGE) to completely replace every genomic occurrence of the amber (UAG) stop codon with the UAA stop codon, freeing the UAG codon for use by the OTS^28^. Together with the deletion of release factor 1 (RF1), this strain dramatically reduces the cellular stress caused by off-target suppression of amber stop codons associated with non-recoded cells and decreases protein truncation caused by premature translation termination in recombinant proteins possessing internal UAG codons^29–31^.

**Figure 1:**
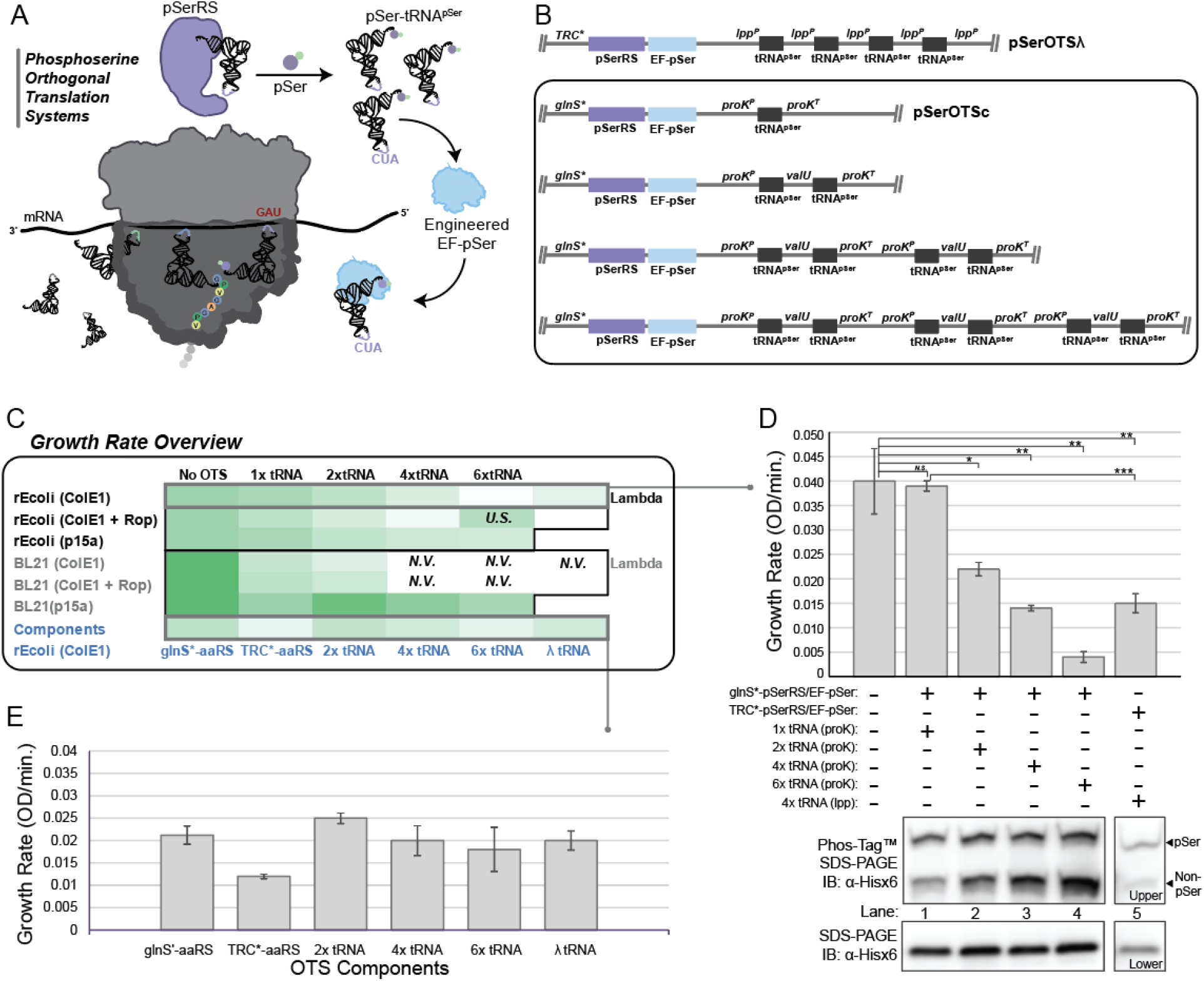
pSerOTS components impact cellular growth and viability. Components of the pSerOTS are illustrated in (**A**). Variations to OTS architecture include the implementation of glnS* (modified *Ec* GlnRS promotor) and the proK (*Ec* tRNA^Pro^) promoter/terminator with valU tRNA linker sequence (*Ec* tRNA^Val^ operon) (**B**) These components are predicted to influence the overall viability of cells harboring OTSs. The effect on the growth of *E. coli* cells expressing various OTSs or components was assessed by determination of kinetic growth parameters through measurement of cell density over time, providing a measure of growth rate. *N.V.* = Not Viable, *U.S.* = Unstable, plasmid easily mutates to reduce tRNA copy (**C**). The impact of tRNA copy number variation on growth and translational fidelity was assessed by comparison of growth rates to relative pSer incorporation into a GFP reporter as determined by Phos-Tag™ SDS-PAGE and immunoblot (**D**). The influence of individual OTS components on growth rate is highlighted in (**E**). All growth parameters reflect the average measurements from three independent replicates with error bars representing 1 *S.D.* Statistical significance of growth rate changes was assessed using an unpaired t-test where * equals p < 0.05, ** equals p < 0.005, *** equals p < 0.0005 and *N.S.* denotes Not Significant.

Outside of strain recoding efforts, the biology of OTS development and specifically the impact on the host organism has been largely overlooked. As implied by the name, OTSs are meant to be orthogonal to host processes and essentially invisible to the cell outside of their use in recombinant protein expression. The introduction of an OTS into the cell adds a foreign element to the make-up of cellular substrate pools. Realistically, the components of the OTS interact with these pools with varying extent. The introduction of nsAAs to the amino acid pool presents host aaRSs with the opportunity to interact with an additional substrate which may lead to misaminoacylation or aaRS inhibition. Likewise, the introduction of a new aaRS:tRNA pair to the cell may challenge the fidelity and efficiency of the host translation machinery which evolved in the absence of this additional substrate^32–35^. Each foreign element introduced into the cell is an opportunity to perturb the balance or fidelity of substrate pools and activate stress responses, like the stringent response. To better understand the impact of OTSs on host physiology, we have explored the extent to which each of our model OTS (pSerOTS) components cause stress within the cell through in-depth profiling of interactions with cellular substrate pools and identification of the potential outcomes of those interactions. Using this information, we construct several pSerOTS variants designed to enhance OTS orthogonality by minimizing the interactions with host substrate pools and decreasing global OTS burden. Overall, we present a framework for the systematic characterization of OTS components and their participation in host cellular processes. This framework can be generally applied to OTS development and should serve as a guide for future OTS design and re-evaluation of existing OTSs with a central focus on enhancing biological tolerance.

## Results

### OTS components impact the growth and viability of host cells

To gain a cursory understanding of the impact OTSs have on the growth and viability of host cells, we constructed pSerOTS derivatives with varying levels of component expression. Derivatives included modulation of expression from the pSerRS:EF-pSer operon, alteration of plasmid origin of replication, and increases in tRNA copy number (**Figure 1B**). We then systematically introduced these OTS variants into divergent lineages of *E. coli*. The first is a fully recoded K-strain MG1655 derivative, rEcoli – experimental phosphoserine (rEcoli^XpS^), and the second is a non-recoded B-strain, BL21 (DE3) routinely used in recombinant protein expression. Each strain carries a deletion for the phosphoserine phosphatase, serB, in the serine biosynthesis pathway to eliminate the conversion pSer to Ser, ensuring an adequate supply of pSer in the amino acid pool^36^. Using these systems, we performed kinetic growth analyses to construct an overview of the potential impact of OTS components on growth and viability **(Figure 1C, Table S2**). Although highly functional, our best pSerOTS variant to date (pSerOTSλ) has been observed to cause growth deficiency and strain instability in some contexts. This presents pSerOTSλ as an ideal model for the present study and therefore used to establish baseline growth parameters for OTS variant benchmarking. In rEcoli^XpS^, inclusion of pSerOTSλ caused a ~2-fold decrease in growth rate when compared to cells lacking the OTS. In addition to a reduction in growth rate, cell size, an indicator of stress in *E. coli*, was increased ~50% in rEcoli^XpS^ cells with pSerOTSλ when compared to rEcoli^XpS^ cells alone (**Figure S1**). Attempts to introduce pSerOTSλ into non-recoded BL21 (DE3) cells were unsuccessful, likely due to the stress imparted by aberrant nonsense suppression events at ORFs terminating in UAG (**Figure 1C**). We next examined OTS variants that were rationally designed using regulatory components successfully deployed in other OTS contexts. The most heavily modified pSerOTS variant placed control of the pSerRS:EF-pSer operon under the constitutive, low level promoter glnS*. A single copy of tRNA^pSer^ was placed under the control of a proK *E. coli* tRNA promoter^37^. This ColE1-based vector was designated pSerOTSc and used as the basis for comparison for OTSs derived from this base lineage (**Figure 1B**). The growth rate of rEcoli^XpS^ cells containing pSerOTSc was nearly identical to that observed in WT cells. This is in stark contrast to the decrease in growth rate observed in cells with pSerOTSλ. pSerOTSc was successfully introduced into non-recoded BL21 (DE3) ΔserB cells, but displayed compromised growth compared to WT BL21 (DE3) ΔserB. Analysis of pSerOTSc variant performance in BL21 (DE3) by Phos-tag™ gel and immunoblot revealed that most BL21 (DE3) strains with OTS variants failed to adequately facilitate pSer incorporation into the E(17)TAG-GFP reporter protein. Successful OTS deployment in BL21 (DE3) depended on the use of a partially genomically recoded strain of BL21 (BL21 (DE3) B-95)^38^, with pSerOTSc exhibiting the best performance overall (**Figure S2**).

tRNA copy number is an important factor in the deployment of a productive OTS^24^. Using pSerOTSc as a base, we created OTS variants with increasing tRNA copy number (**Figure 1B**). We introduced these OTS variants into both rEcoli^XpS^ and BL21 (DE3) ΔserB. In rEcoli^XpS^, we observed a dose dependent decrease in growth rate as tRNA copy number was increased. The same trend held for BL21 (DE3) ΔserB, however cells became inviable with OTS variants having greater than two copies of tRNA. Similar results were obtained using OTS variants with identical OTS components but altered to include the Rop protein to mediate a decrease in tRNA copy number. The only exception to OTS viability extending beyond the inclusion of 2x tRNA in BL21 (DE3) ΔserB was when the origin of replication on the OTS plasmid was switched to lower copy p15a. Both rEcoli^XpS^ and BL21 (DE3) cells harboring OTS variants with p15a origins were viable and generally tolerated increases in tRNA copy number (**Figure 1C-D**).

To narrow down the potential contribution of each OTS component to the observed growth differences, we constructed variants expressing only individual OTS components. The first component we isolated for expression in the host cell was the aaRS. As a starting point, we modified the existing pSerOTSλ OTS to excise the tRNA and EF-pSer components, leaving pSerRS under the control of the modified TRC promoter (TRC*) which provided constitutive, high level of pSerRS. For comparison, we constructed an additional vector expressing pSerRS under control of glnS*. The growth of component OTS vectors was only measured in rEcoli^XpS^ to reduce confounding growth defects due to codon suppression. In comparison to *E. coli* lacking an OTS (WT), cells expressing glnS*-pSerRS showed a slight growth defect, while cells expressing TRC*-pSerRS displayed a significant decrease in growth (**Figure 1E**). We constructed tRNA only plasmids by removing the pSerRS:EF-pSer operon from vectors illustrated in Figure 1B. For the pSerOTSc derived tRNA variant plasmid, a similar tRNA copy number dependent decrease in growth rate to that of the counterparts with pSerRS:EF-pSer was observed. In contrast, the pSerOTSλ derived tRNA only plasmid (4x tRNA driven by five independent lpp promoters) displayed improved growth performance when compared to the pSerRS only and full pSerOTSλ variants. Altogether, analysis of the impact of full and individual OTS components on growth suggests that both pSerRS level and tRNA copy number are important factors influencing cellular viability.

While host tolerance is essential to the successful deployment of an OTS, function and fidelity are equally as important. To assess the performance of the newly constructed pSerOTSc lineage against that established by the progenitor pSerOTSλ, we expressed E(17)TAG-GFP reporter alongside each OTS variant. GFP containing pSer was separated from non-pSer containing GFP using a Phos-tag™ SDS-PAGE gel-shift assay visualized by immunoblot against a C-terminal 6xHis epitope tag. GFP reporter expression facilitated by pSerOTSλ was decreased when compared to pSerOTSc variants. Interestingly, the tRNA copy number dependent decrease in cellular viability was inversely proportional to the amount of non-pSer containing GFP reporter (**Figure 1D**). This observation suggests that tRNA^pSer^ misaminoacylation may play a role observed decrease in viability. Taken together, the preliminary growth analysis and corresponding reporter expression data indicate that each component of the OTS, and it’s relative level of expression, is critically important to both host tolerance and OTS fidelity.

### High-level orthogonal aaRS expression redirects cellular resources and decreases aa-tRNA pool fidelity

When an OTS is introduced into a cell, the cell must redirect transcriptional and translational resources towards the production of OTS components. Typically an OTS is used in conjunction with additional vectors expressing recombinant protein targeted by the OTS. Thus, the commitment of resources to OTS expression needs to be carefully weighed against the expression of the target recombinant protein. Even then, the possibility remains that the impact of the OTS on cellular physiology may prevent efficient expression of either system, as evident by the growth defects and variable reporter expression levels observed in Figure 1. To better understand the implications of OTS expression on the host cell, we performed in-depth proteomic analysis of rEcoli^XpS^ in different pSerOTS contexts with particular emphasis on the expression of pSerRS and its impact on host physiology. To measure pSerRS levels, rEcoli^XpS^ cells were transformed with pSerOTSλ, pSerOTSc, or Rop/p15a versions of pSerOTSc and grown to mid-log in nutritionally replete medium. Total protein was extracted from each condition, subjected to proteolytic digest, and quantified using a shotgun proteomics approach. Analysis of the proteomic data yielded rank-ordered lists based on relative abundance of individual *E. coli* and OTS proteins. pSerRS and native *E. coli* aaRSs were plotted for each experimental condition to illustrate the abundance of pSerRS compared to the rest of the aaRS pool within the context of the *E. coli* proteome (**Figure 2A**). Surprisingly, the level of pSerRS from pSerOTSλ (top 10 most abundant) indicated it was one of the most abundant proteins in the cell; substantially higher than any native aaRS level which typically rank 300-400. Beyond the extreme metabolic demand of maintaining high-level expression during log-phase growth, the abundance of pSerRS is likely to challenge the cell by disrupting the binding equilibrium within amino acid and tRNA substrate pools. This has previously been demonstrated to decrease enzymatic selectivity resulting in mistranslation events in both *E. coli* and *S. cerevisiae^39,40^.* Expression of pSerRS from pSerOTSc fell within the range of expression of native aaRSs. Overall, pSerRS expression from the pSerOTSc copy number variants decreased with plasmid copy number, as expected (**Figure 2A-B**).

**Figure 2:**
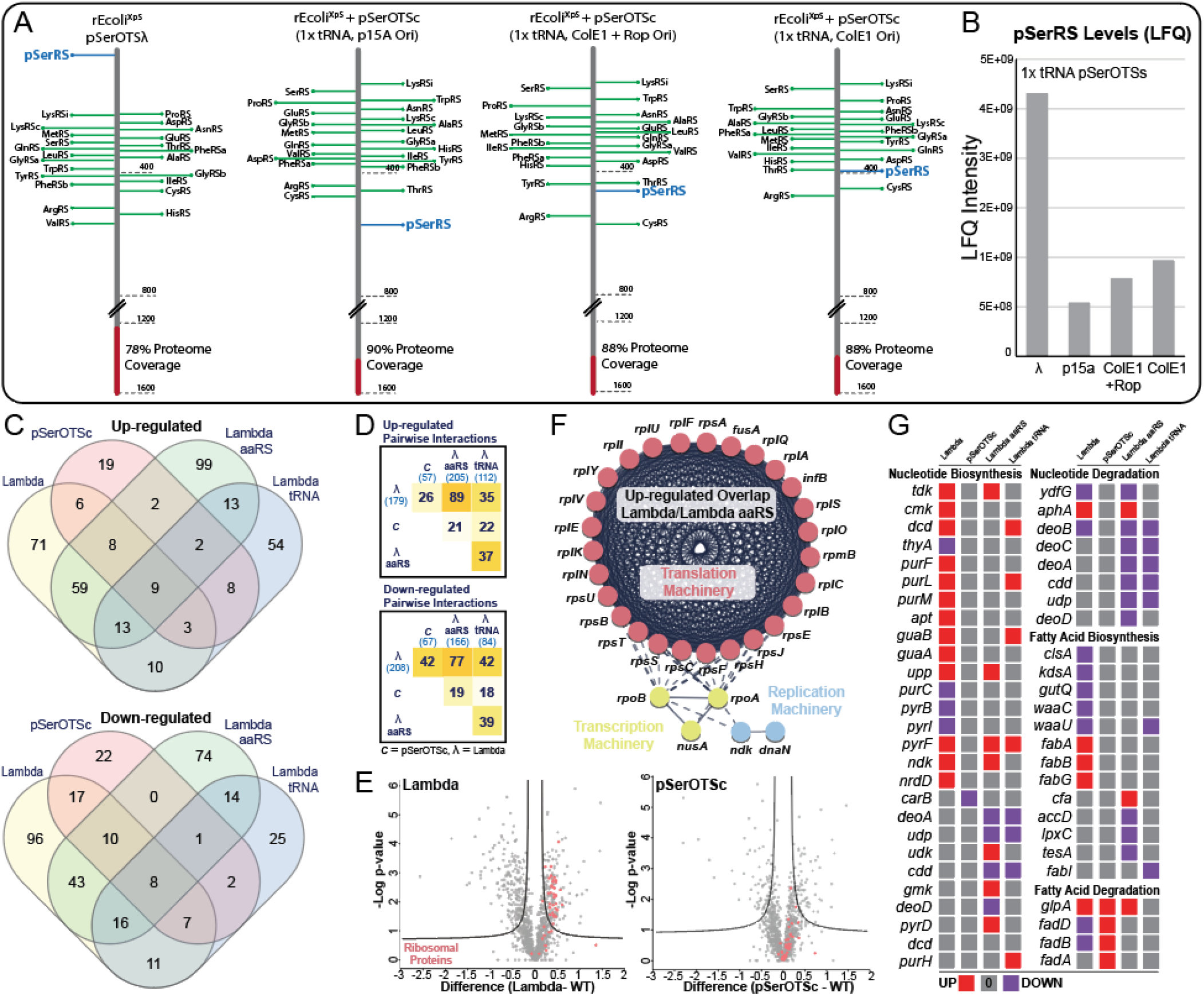
pSerRS expression levels modulate cellular fitness and stress response. The level of pSerRS protein in cells containing various OTS constructs was measured by mass spectrometry and the position of pSerRS (blue line) relative to all proteins detected in the proteome is displayed in the context of native aaRS levels (green lines) (**A**). Total pSerRS levels were determined by label-free quantification and graphed as a function of OTS variant (**B**). rEcoli^XpS^ cells expressing complete or partial OTS constructs were subjected to proteomic analysis. The overlap of significantly upregulated and downregulated proteins is summarized in (**C**). Pairwise overlap across experimental conditions is highlighted in (**D**). Alteration of the proteomes from cells containing pSerOTSλ and pSerOTSc are illustrated as volcano plots by plotting functions of the p-value against differences in proteome composition compared to WT cells without an OTS. Significant proteins are indicated as those outside the asymptote (black lines) with the expression of ribosomal proteins highlighted in pink (**E**). The overlap in upregulated proteins from proteomes impacted by pSerOTSλ and Lambda aaRS alone was input into stringDB to identify statistically significant pathway enrichment and grouped using k-means clustering algorithm. Known protein-protein interactions were enriched for upregulation in translational (pink), transcriptional (yellow), and replication (blue) machinery (**F**). Pathway enrichment analysis was conducted for strains harboring OTS variants using Pathway Tools and to illustrate the regulation of the highly impacted pathway components with up-regulated proteins in red, down-regulated proteins in purple, and proteins with no change compared to WT in grey. Enrichment cutoffs were set to a statistically significant differential expression score of 0.1 (**G**). All proteomes were quantified in triplicate and analyzed in Perseus using t-test and volcano plot functions to obtain statistically significant proteomic deviations.

Comprehensive proteomic analysis also provides a snapshot of active cellular processes which allow us to deduce protein networks and infer deleterious responses to OTS challenge. To further understand the effects of each OTS component on the host cell, we expanded our proteomic analysis to include rEcoli^XpS^ expressing only the tRNA (Lambda tRNA) or pSerRS (Lambda aaRS) components of pSerOTSλ (Lambda). Together with the full OTS proteomic data, we can attribute proteomic responses to each OTS constituent. We compiled the rank ordered protein abundance lists from each condition obtained in triplicate and performed statistical analysis to determine the proteins whose expression deviates significantly from the control rEcoli^XpS^ proteome devoid of OTS influence. The lists of proteins with significantly up-regulated and down-regulated protein expression compared to the rEcoli^XpS^ control were cross referenced to identify overlap between each experimental condition (**Figure 2C**). Pairwise comparison of significant proteins across the sample conditions highlights the substantial difference in the total number of significantly divergent proteins between pSerOTSc and pSerOTSλ-derived OTSs. pSerOTSc had 57 up-regulated proteins and 67 down-regulated proteins compared to the rEcoli^XpS^ control, while Lambda had 179 up-regulated and 208 down-regulated proteins. Similar levels were observed in the Lambda aaRS only proteome (**Figure 2D**). The observed increase in dis-regulated proteins relative to the pSerOTSc proteome underscores the extent to which Lambda perturbs the proteome of the host cell. These data also capture the extent that a host proteome differs with an OTS, in general, which further highlights the host cell burden.

To further understand OTS-mediated changes to the proteome, we constructed volcano plots comprised of every up- and down-regulated protein expressed in Lambda and pSerOTSc relative to the rEcoli^XpS^ control. Points which lie on the right side of each plot represent up-regulated proteins, while those above the curved boundary line are significant. We hypothesized that constant demand from pSerRS expression on pSerOTSλ might require an increase in translational capacity within the cell and examined levels of ribosomal protein subunits (highlighted in red) as a proxy for translational demand. As illustrated, Lambda containing cells display a significant increase in the levels of ribosomal proteins compared to cells containing pSerOTSc (**Figure 2E**). In congruence with the highlighted difference in ribosomal protein expression, the complete overview of significant proteomic changes overlaid on the *E. coli* genome illustrates generalized effect of high-level pSerRS expression on the proteome (**Figures S3-S4**). These data clearly illustrate the intense translational demand imparted by pSerRS expression from pSerOTSλ which results in substantial redirection of cellular resources towards the production of ribosomes. This observation may also help to explain the perceived reduction in recombinant reporter expression from these cells when compared to cells utilizing a more conservative pSerOTSc variant. Unique overlap of up-regulated proteins between Lambda and the Lambda aaRS variant confirms that the upregulation of ribosomal proteins is the result of high-level pSerRS expression. Further analysis of these sample groups revealed up-regulation of RNA polymerase subunits, likely required to support high level transcription of pSerRS (**Figure 2F**). Additional pathway analysis of the proteomic results from the four sample groups (Lambda, pSerOTSc, Lambda tRNA, and Lambda aaRS) revealed OTS-composition dependent changes in metabolic processes within the host cell. In addition to the translational demands of Lambda, the cell also up-regulates the expression of proteins involved in nucleotide biosynthesis, while simultaneously down-regulating proteins involved in nucleotide degradation (**Figure 2G**). This metabolic reprogramming may be required to support the combined transcriptional demand of the pSerRS:EF-pSer operon and tRNA cassette on pSerOTSλ (**Figure 2G**). Lambda also elicits a unique cellular response through up-regulation of proteins involved in fatty acid biosynthesis and concomitant decrease in fatty acid degradation proteins. The cause of this metabolic reprogramming event is less readily apparent and may be related to carbon-mediated short chain fatty acid stringent response activation or perturbation of regulatory proteins^41,42^ (**Figure 2G, Figure S5**). Unique metabolic reprogramming in the tRNA only OTS sample centered around up-regulation of amino acid biosynthesis, a hallmark of stringent response activation due to an increase in deacylated tRNA pools^43^ (**Figure S6**). While pSerOTSλ, and OTSs derived from pSerOTSλ, severely perturbed proteomic homeostasis, relatively few alterations to the proteome were observed during expression of the pSerOTSc system that correspond well with OTS-mediated growth characteristics. Overall, these data reinforce the notion that fine-tuning OTS component expression based on in-depth proteomic analysis can minimize the impact of an OTS on the host cell.

From amino acids to tRNAs, aaRSs interact with a wide range of cellular substrate pools. Extensive literature on aaRS enzymology both *in vitro* and *in vivo* has illustrated the potential for high level expression of aaRSs to disrupt the kinetic environment leading to competition between aaRSs for substrate binding often resulting in misacylation events^39,40^. Even so, misaminoacylation of the native tRNA pool due to the introduction of an orthogonal aaRS has never been systematically profiled. While interactions between pSerRS and native tRNAs have been explored in a limited context *in vitro*, numerous proofreading mechanisms exist in the cell post-aminoacylation which may prevent misincorporation of pSer into the proteome^*44–47*^. To investigate whether high-level expression of pSerRS facilitates pSer misincorporation *in vivo*, we leveraged our recently developed Mass Spectrometry Reporters for Exact Amino Acid Decoding (MS-READ) reporters. The flexible design of this reporter protein enabled us to investigate misincorporation events for various native codons at the guest position of the reporter peptide^48^. Codons for the amino acids Gly, Ser, and Thr, were initially chosen for their similarities in tRNA recognition elements (outlined below). MS-READ analysis revealed that the majority of the incorporation events were faithful to the amino acid coded at the guest position. Only the Gly reporter had a measurable level of pSer misincorporation (**Figure 3A, Figure S7**). Careful examination of the tRNA recognition elements used by aaRSs for tRNA selection shared between tRNA^Gly^ and tRNA^pSer^ identified significant overlap in major tRNA^pSer^ recognition elements in the primary sequence of tRNA^Gly^ (**Figure 3B**). This suggests a mechanism for misaminoacylation of pSer onto tRNA^Gly^ due to tRNA misrecognition and provides *in vivo* evidence of *in vitro* observations^18,49^. We next re-examined our proteomic data for pSer misincorporation events at native Gly positions in host proteins. Without phospho-enrichment strategies, we found a total of 33 unique peptides with pSer misincorporation at native Gly codons (**Table S3**). An example of one such event at Gly110 of groL is shown in **Figure 3C**. The misincorporation of pSer at Gly codons is an important example of native proteome damage caused by an orthogonal aaRS and highlights importance of monitoring and tuning aaRS expression during OTS development.

**Figure 3:**
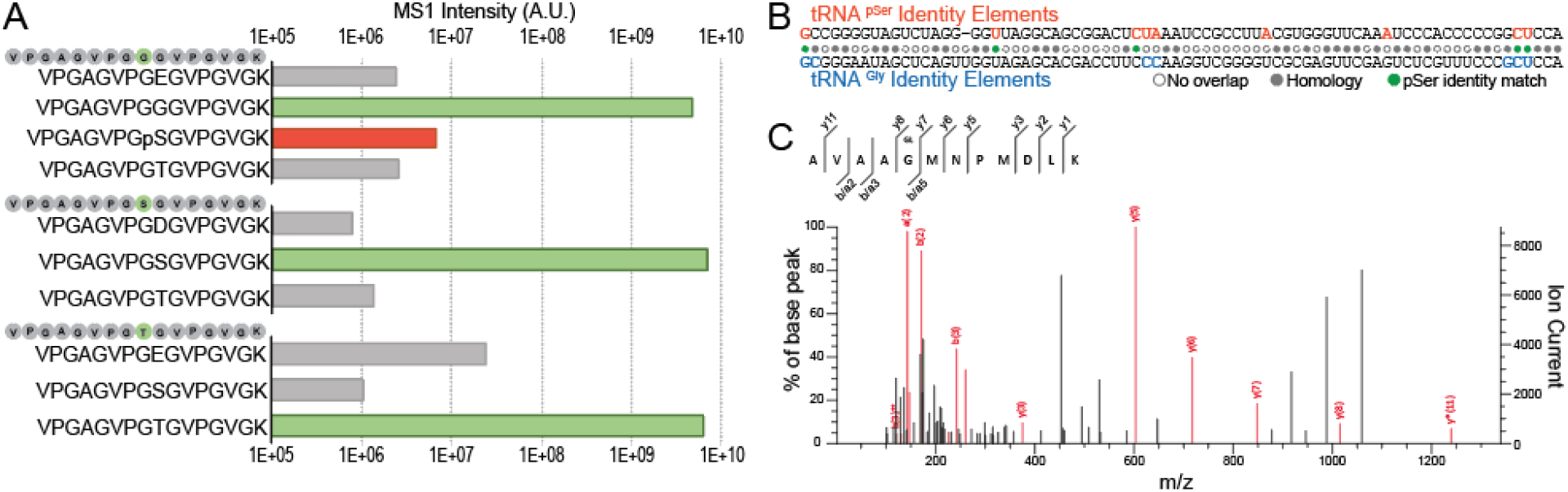
High level pSerRS expression results in proteome-wide pSer misincorporation. Decoding fidelity at Gly, Ser, and Thr codons was assessed by monitoring site-specific amino acid incorporation. Amino acids incorporated at the indicated reporter position (green circle) were quantified by mass spectrometry and incorporation events were graphed to display faithful decoding (green bars), misincorporation (grey bars), and pSer misincorporation (red bars) as a function of MS1 precursor ion intensity (**A**). tRNA identity elements for *E. coli* tRNA^Gly^ (blue) and tRNA^pSer^ (orange) were highlighted on the tRNA primary sequence and aligned to identify primary sequence overlap (grey circles) and pSerRS recognition elements common to both tRNAs (green circles) (**B**). The proteomes from pSerOTSλ and Lambda aaRS only cells were searched for pSer misincorporation using custom modification parameters in Mascot. The MS2 spectrum and sequence for example groL peptide (m/z 664.293 M^++^) found in all samples is presented in (**C**).

### Balance of orthogonal tRNA in the tRNA pool is essential to translational fidelity

Specific OTS components may affect recombinant protein quality in addition to yield. We noticed that increasing the copy number of the suppressor tRNA while decreasing the levels of pSerRS led to an increase in non-phospho recombinant protein (**Figure 1D**). Two major sources of tRNA-mediated reductions in OTS fidelity are near-cognate suppression by host tRNAs and misaminoacylation of tRNA^pSer^ by native aaRSs^50–52^. This occurs by an orthogonal tRNA disrupting the composition of the native tRNA pool and interfering with native aminoacylation kinetics. To better understand the implications of orthogonal tRNA introduction to the host cell, we thoroughly characterized the interactions between tRNA^pSer^ and host physiology with particular emphasis on translational fidelity.

The suppression of stop codons occurs naturally in protein translation by ribosomal read-through using near-cognate aa-tRNAs and, more spontaneously, by mutations to the anticodon region of a tRNA resulting in a suppressor tRNA^53,54^. In their native context, many of these read-through events arise at random, but some serve regulatory roles or facilitate priming the cell for enhanced stress tolerance through decreased translational fidelity resulting in erroneous protein synthesis and misfolding^55–58^. In rEcoli^XpS^, a strain fully recoded to remove all UAG stop codons, the cells access to the UAG codon has been limited to instances in recombinantly expressed proteins. To assess the interaction between the recoded cells native aa-tRNA pool and artificially restricted codon, we measured the rates of amino acid misincorporation by near-cognate suppression using MS-READ. rEcoli^XpS^ cells were transformed with an MS-READ reporter plasmid with a UAG codon in the guest position of the reporter peptide. Amino acid incorporation events were observed in the absence of an OTS. Analysis of the purified reporter revealed that the majority of the misincorporation events were Phe and Gln, with moderate levels of Tyr and low levels of Lys and Asn also observed (**Figure 4A**). The incorporation of Gln, Tyr, and Lys has been previously observed as UAG near-cognate suppression events, however the relatively high level of Phe incorporation is unexpected and may be an artifact unique to rEcoli^XpS^ cells^59^. As a whole, these data establish the fundamental baseline for native near-cognate and non-cognate suppression events in rEcoli^XpS^ cells at UAG codons in the absence of OTS components.

**Figure 4:**
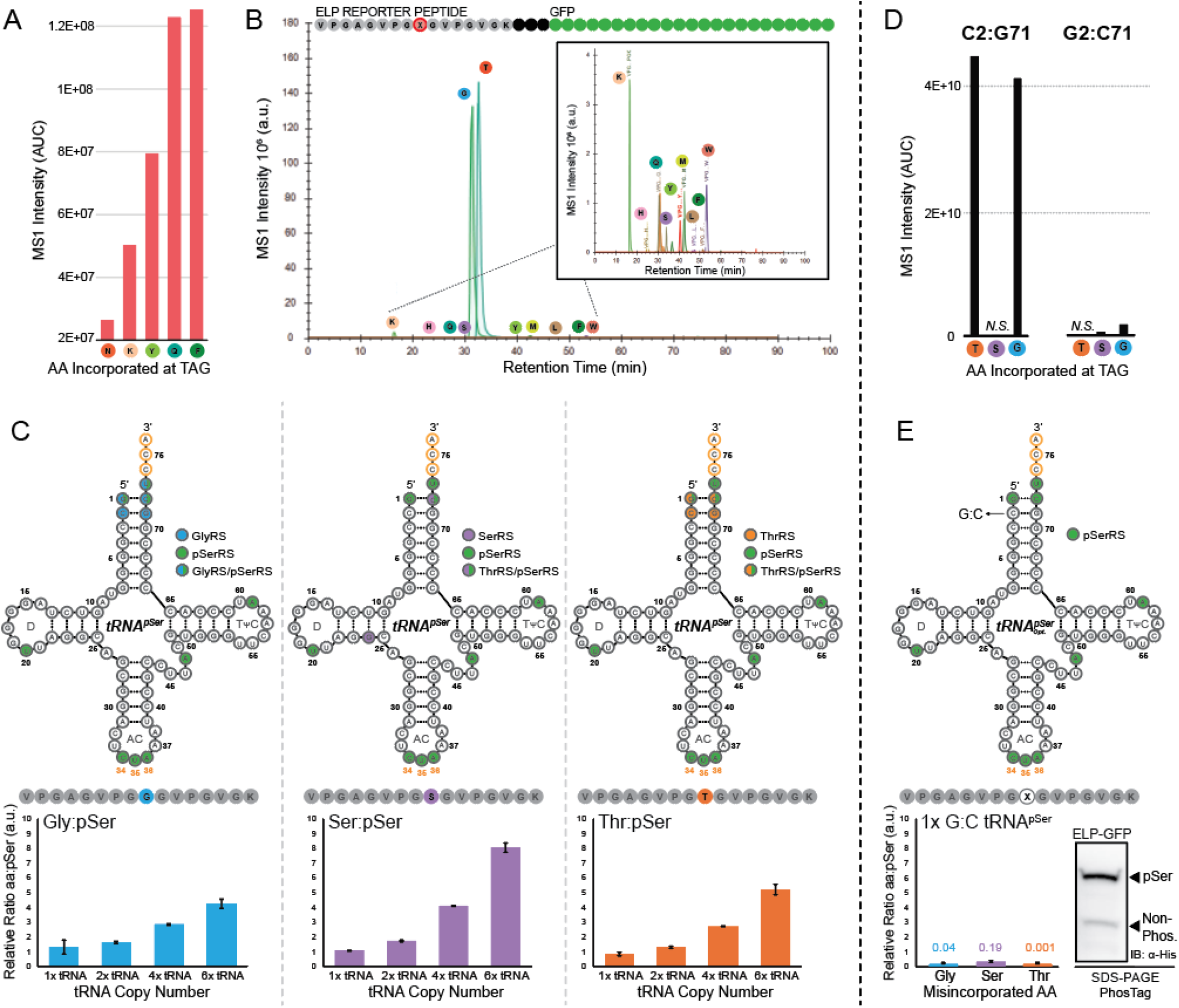
tRNA identity overlap enables concentration-dependent misaminoacylation. Amino acid incorporation was monitored at a UAG-containing MS-READ reporter in the absence of an OTS by mass spectrometry. Incorporation was analyzed using Skyline and MS1 precursor ion intensities (area under the curve) was graphed as a function of amino acid identity (**A**). Misaminoacylation by native aaRSs was identified by following incorporation events in a UAG-MS-READ reporter expressed in the presence of orthogonal tRNA^pSer^. Incorporation was quantified using Skyline and all incorporation events were displayed by MS1 intensity as a function of retention time (**B**). *E. coli* tRNA identity elements were overlaid onto the primary tRNA^pSer^ sequence. The effect of tRNA copy number increase on misaminoacylation was followed by mass spectrometry using a UAG-MS-READ reporter in the presence of OTSs with increasing tRNA copy number. Incorporation events from three independent samples were quantified using Skyline and graphed relative to the level of pSer incorporation within the same sample (**C**). The effect of tRNA modification C2:G71 on aminoacylation fidelity was assessed by measuring amino acid incorporation in the at UAG-MS-READ reporter expressed in the presence of modified tRNA^pSer^ only. Incorporation was analyzed using Skyline and MS1 precursor ion intensities (area under the curve) was graphed as a function of amino acid identity (**D**). The effect of tRNA modification on misaminoacylation was followed by mass spectrometry using a UAG-MS-READ reporter in the presence of modified tRNA. Incorporation events from three independent samples were quantified using Skyline and graphed relative to the level of pSer incorporation within the same sample (**E**).

Orthogonal tRNAs challenge cells with foreign, yet highly similar substrates for molecular interactions. The limited chemical variability afforded tRNAs by their four base composition creates issues with tRNA recognition element overlap across isoacceptor groups^60,61^. To identify the elements of orthogonal tRNA^pSer^ which enable a steady state pool of misaminoacylated tRNA and results in misincorporation, we again used the MS-READ reporter expressed in rEcoli^XpS^ cells alongside a vector containing two copies of tRNA^pSer^. Analysis of the purified reporter protein revealed high-level misincorporation of Gly and Thr compared to background misincorporation events (**Figure 4B**). These misincorporation events are 100-fold greater than near-cognate suppression events identified in the reporter only sample (**Figure 4A**), suggesting that these events are more likely the result of misaminoacylation of tRNA^pSer^ with Gly and Thr rather than nonsense suppression mediated by native tRNA. The identity elements that govern the interactions of both glycyl-tRNA synthetase (GlyRS) and threonyl-tRNA synthetase (ThrRS) have been extensively characterized in prior work^49,62,63^. Using these data, we mapped the overlap of tRNA recognition elements onto the primary sequence of tRNA^pSer^ (**Figure 4C**). For both tRNA^Gly^ and tRNA^Thr^, major recognition elements are found in the acceptor stem of tRNA^pSer^ at G1:C72 and C2:G71 and are compatible with the discriminator base U73 used for tRNA^pSer^ recognition^18^. The inclusion of these identity elements in the primary sequence of the orthogonal tRNA^pSer^ demonstrates how the introduction of orthogonal tRNAs into the tRNA pool can lead to aberrant interactions and, in this case, results in misrecognition and misaminoacylation of tRNA^pSer^ by Gly- and ThrRS. Orthogonal tRNA misaminoacylation profiling, in general, should be considered an essential component of OTS development as it provides key insights in tRNA orthogonality and informs changes that could lead to enhanced biological tolerance.

After profiling the interactions of orthogonal aaRS and tRNA components with the host cell, we focused our analysis on host interactions with a complete OTS. Variants of pSerOTSc with 1x, 2x, 4x, or 6x tRNA^pSer^ were co-transformed into rEcoli^XpS^ cells with the MS-READ reporter to monitor decoding of the UAG codon. The results revealed that the tRNA-dependent increase in non-pSer containing reporter protein observed in Figure 1D is likely the result of increasing tRNA misaminoacylation. At low tRNA copy number (e.g. 1x tRNA), misaminoacylation is relatively low, but as the pool of orthogonal tRNA is expanded in cells, a dose-dependent increase in misaminoacylation by native aminoacyl-tRNA synthetases leads to a marked increase in the relative incorporation rates of Gly and Thr relative to pSer (**Figure 4C**). Interestingly, high level Ser misincorporation was also observed alongside Gly and Thr misincorporation. Ser misincorporation required both orthogonal aaRS and tRNA expression and was absent in isolation of either component, ruling out misaminoacylation of tRNA^pSer^ by seryl-tRNA synthetase (SerRS) and near cognate suppression. Mapping of SerRS recognition elements onto tRNA^pSer^ primary sequence highlights the low overlap of SerRS recognition elements, providing further evidence against native SerRS interaction with orthogonal tRNA^pSer^ (**Figure 4C**)^64^. Ruling out misaminoacylation by SerRS as the cause of Ser misincorporation leaves either pSer de-phosphorylation post-incorporation, or misaminoacylation of tRNA^pSer^ with Ser as possible mechanisms. While de-phosphorylation of pSer may contribute to the Ser levels, the tRNA-dependent increase in the relative ratio of Ser:pSer points towards a specific interaction between pSerRS and tRNA^pSer^ which results in Ser misaminoacylation (**Figure 4C**). Taken together, these results illustrate the potential for OTS components to interact with host translational machinery, resulting in perturbation of substrate pools and relatively high-level mistranslation events.

Using the information gained through monitoring host interactions with pSerOTS, we were able to identify a modification to the OTS which may mitigate a large proportion of pSerOTS-mediated non-cognate interactions within the cell. The identification of tRNA recognition element overlap and the seemingly heavy dependence on tRNA interactions for misincorporation events led to the targeting of tRNA^pSer^ for modification. While the G1:C72 pair in the accepter stem is essential for pSerRS recognition of tRNA^pSer^, the C2:G71 base pairing on tRNA^pSer^ is only recognized by GlyRS and ThrRS, making it an excellent target for modification to reduce off target interaction. To maintain the GC-content of tRNA^pSer^, we swapped the base pair to create a G2:C71 variant of tRNA^pSer^ 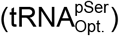. Alteration of this specific tRNA base pair has been targeted as a means to decrease GlyRS recognition previously, but with a different OTS and without thorough characterization of the interactions between the sequence-modified tRNA and the cell^19^. To determine whether 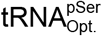 would reduce Gly and Thr misincorporation rates, we created an OTS variant possessing a single copy of 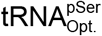 and monitored misaminoacylation mediated misincorporation events using the UAG MS-READ reporter. As we established above, providing the cell with the sequence-modified tRNA in the absence of the orthogonal aaRS is the most stringent and challenging test for interaction fidelity between the orthogonal tRNA and host aaRSs. Under these conditions, we observed a substantial reduction in Gly misincorporation (~100-fold) when compared to WT tRNA^pSer^ under the same experimental conditions. Amazingly, the interactions between tRNA^pSer^ and ThrRS was practically ablated, with any Thr misincorporation events falling below the limit of detection for this experiment (**Figure 4D**).

With these promising results, we constructed a variant of pSerOTSc which included one copy of the sequence-modified tRNA (pSerOTSc*). pSerOTSc* was co-transformed with a UAG MS-READ reporter for expression in rEcoli^XpS^. In comparison to the OTS with WT tRNA^pSer^, the relative incorporation rate of Gly:pSer for the OTS with 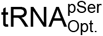. was reduced at least 30-fold, while the relative incorporation rate of Thr:pSer was reduced a remarkable 1000-fold (**Figure 4E**). At the same time, a large decrease in Ser:pSer relative incorporation rates were also observed, suggesting that the sequence modification made to the orthogonal tRNA may have reduced Ser misaminoacylation (**Figure 4E**). A Phos-tag™ assay reflected the increase in recombinant protein purity quantified in the MS experiment (**Figure 4E**). As a whole, these results illustrate the need to carefully probe the interactions between OTS components and the host cell. Orthogonal tRNA, in particular, is centrally positioned as an intermediary of protein synthesis and interacts with a wide range of cellular substrate pools. The substantial improvement in pSerOTS fidelity through tRNA copy number modulation and sequence modification is a prime example of the power of data-driven OTS design and its ability to reduce host toxicity.

### Tuning of OTS components to enhance orthogonality improves protein yield and purity

Though careful design and systematic testing of components during OTS assembly and evolution can help to mitigate deleterious outcomes, OTS function should ultimately be judged by its ability to produce a functional target protein of interest. To test the performance of pSerOTSc **(Figure 5A**), we expressed full-length doubly phosphorylated human MEK1. In human cells, MEK1 is activated by phosphorylation at residues S218 and S222 in the activation loop^65^. We have previously expressed variants of phosph-MEK1 using pSerOTSλ which creates an ideal frame of reference for OTS performance^18^. The MEK1 (2x UAG) expression vector was co-transformed with an OTS plasmid (pSerOTSλ or pSerOTSc) or a plasmid carrying a suppressor tRNA^Ser^ (supD) as a negative control into rEcoli^XpS^ cells or into a serB competent rEcoli^XpS^. Following expression, crude protein lysate was separated by SDS-PAGE and MEK1 phosphorylation status was assessed using a phospho-specific antibody which recognizes MEK1 pS218/pS222. Lanes 2 and 3 (upper) show phospho-MEK1 expression using pSerOTSλ and pSerOTSc, respectively, in rEcoli cells. The presence of serB in these cells tests the limits of pSerOTS efficiency and selectivity by decreasing the intracellular pool of pSer relative to near-cognate amino acids. Under these conditions, pSerOTSc (Lane 3) is able to robustly outperform pSerOTSλ (Lane 2) when compared to total and phospho-MEK1 expression. A similar trend is observed in rEcoli^XpS^ cells where the pool of intracellular pSer has been restored (Lanes 4-5, **Figure 5B**). To further examine performance in the context of phospho-specificity, we expressed a split-mCherry fluorescent reporter in the presence of the pSerOTSc tRNA copy number variants. This reporter reconstitutes fluorescence when one half of mCherry fused to the human phospho-binding protein 14-3-3β interacts with its phosphorylated peptide substrate fused to the second half of the mCherry reporter, thereby allowing productive interactions to be observed with single cell resolution via flow cytometry^66^. Following the trend observed in Figure 1D, increase in tRNA copy number resulted in a decrease in relative pSer incorporation and, in turn, a decrease in productive protein-protein interactions. When compared to pSerOTSλ, pSerOTSc with 1x tRNA displayed no discernable difference in fluorescence shift compared to the no OTS negative control sample indicating that pSerOTSc is able to robustly facilitate phospho-dependent protein-protein interactions *in vivo* (**Figure S8**). Overall, the changes made to the OTS architecture resulted in an OTS variant which is less toxic to the host and allows for redirection of cellular resources to facilitate phosphorylation dependent protein-protein interactions and the faithful production of a doubly phosphorylated recombinant protein.

**Figure 5:**
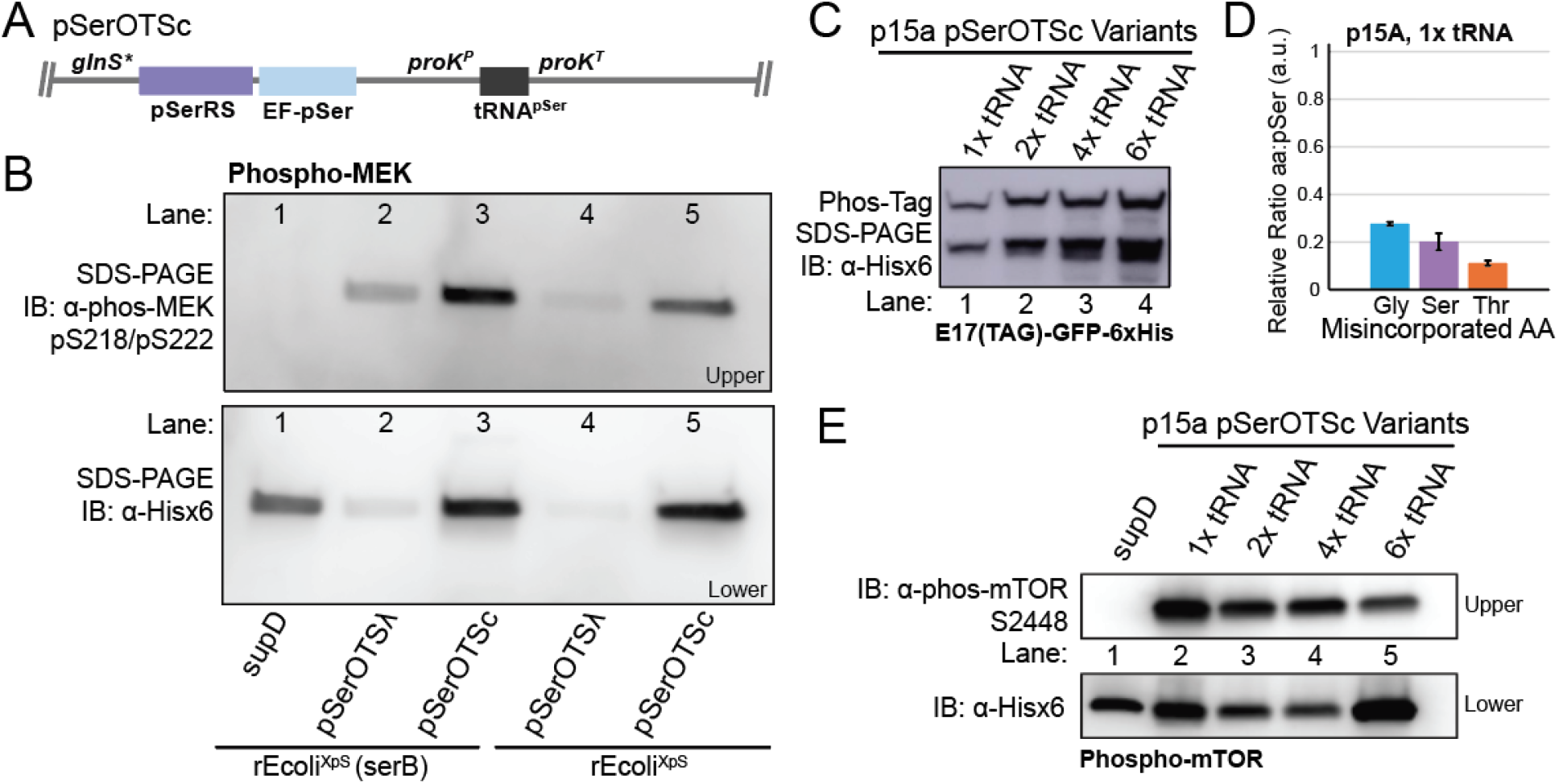
Examination of OTS variant performance. Architecture of optimized pSerOTSc (**A**). pSerOTSc performance relative to the progenitor pSerOTSλ was assessed in both serB competent (Lanes 1-3) and serB deficient (Lanes4-5) strains by expression of recombinant MEK1 containing two sites for pSer incorporation and visualized by immunoblot using α-phospho-MEK1 S218/S222 (**B**). The effect of plasmid copy number variation on OTS fidelity was examined by Phos-tag™ gel (**C**). The effect of p15a OTS variants on misaminoacylation was followed by mass spectrometry using a UAG-MS-READ reporter in the presence of modified tRNA. Incorporation events from three independent samples were quantified using Skyline and graphed relative to the level of pSer incorporation within the same sample (**D**). The performance of pSerOTSc p15a variants were assessed by expression of a recombinant human mTOR fragment containing a single site for pSer incorporation and visualized by immunoblot using α-phospho-mTOR S2448 (**E**).

Having established that pSerOTSc works robustly with a ColE1 ORI (~30-40 copies/cell), we next sought to establish whether decreasing the plasmid copy number by changing the ORI to p15a (~10-20 copies/cell) had any impact on overall OTS performance. pSerOTSc p15a variants with increasing tRNA copy number (1x-, 2x-, 4x-, and 6x-tRNA) were co-transformed with the E(17)TAG-GFP reporter into rEcoli^XpS^ cells and analyzed by Phos-tag™ assay. With p15a ORIs, the OTS variants display the similar trend of increasing misaminoacylation concomitant with an increase in tRNA abundance observed in the pSerOTSc ColE1 architecture (**Figure 5C**).

To more closely examine the impact of copy number decreases on OTS fidelity, the 1x tRNA pSerOTSc p15a variant was co-transformed with the UAG MS-READ reporter into rEcoli^XpS^ cells. The same three misincorporation events observed in the ColE1-based OTS (Gly, Thr, and Ser) were also observed in the p15a-based OTS. Misincorporation rates relative to pSer incorporation, however, were between 3-5 fold lower than those observed in the ColE1 variant (**Figure 5D**). The additional increase to OTS fidelity when plasmid copy number is decreased is reflective of the decrease in orthogonal tRNA in the total tRNA pool, lowering the concentration well below the K_m_ for native aaRSs. As a final test of p15a OTS variant performance, we examined its ability to support expression of a recombinant protein fragment containing human phospho-mTOR. The pSerOTSc p15a variants, along with supD as a negative control, were separately transformed into rEcoli^XpS^ cells alongside the UAG-containing mTOR expression vector. Cell lysate was separated by SDS-PAGE and visualized by immunoblot using an antibody specific to the phosphorylated version of mTOR, and then re-probed using an antibody specific to the 6xHis c-terminal epitope tag to visualize total mTOR-fragment expression^66^. As expected, the negative control supD expression (Lane 1, upper) failed to illicit a reaction with the phospho-mTOR antibody, but the presence of bands in Lanes 2-5 (upper) indicates all pSerOTSc p15a variants facilitate site-specific expression of the phospho-mTOR fragment (**Figure 5E**). Total expression levels varied across experimental conditions (Lanes 1-5, lower), but the effect of orthogonal tRNA overexpression on recombinant protein purity follows previously observed trends. The 6x tRNA OTS, for example, is robustly expressed (Lane 6, lower), but the proportion of phospho-mTOR production (Lane 6, upper) is significantly lower when compared to the 1x tRNA OTS counterpart (Lane1) (**Figure 5E**).

## Discussion

### Understanding the role of the host in OTS performance

Hundreds of OTSs have been established to facilitate the incorporation of vast collection of nsAA. These OTSs are routinely deployed to characterize protein:substrate interactions, further our understanding of enzyme function, and create synthetically modified proteins with industrial applications. Though OTSs have been established across all domains of life, most are still implemented in *E.coli*. Despite this fact, there remains a huge gap in our understanding of the interactions between OTSs and the host cell and the resulting impact of these interactions on target protein expression. Using pSerOTS as an example, this study provides the first systematic characterization of the global effects of OTS introduction to host *E. coli* cells. It should come as no surprise that, in most cases, *E. coli* acts as more than a vehicle for OTS-mediated protein expression by exerting influence on transcriptional and translational efficiency through stress responses and resource allocation.

The implementation of OTSs for most recombinant protein expression is performed in typical protein expression strains (e.g. BL21) modified to reduce proteolytic degradation of proteins to enhance general recombinant protein yield. While amenable to overall protein expression, this feature dramatically reduces the mechanisms available to the cell to counteract the inevitable nonsense suppression events caused by orthogonal suppressor tRNA and premature translation termination, thus greatly reducing yield in the presence of an OTS. The development of genetically recoded organisms which reduce or eliminate codons targeted by an OTS has provided an effective means to bypass the detrimental effects of off target stop codon suppression. Even in this context, the potential for deleterious interactions with the host cell remains through a break-down in OTS orthogonality.

OTS orthogonality is often regarded as a static condition. However, much like the host cell itself, this simplification is complicated by the dynamic nature of the cell’s response to both external stimuli and internally by the basal demands of physiological processes. For the first time, our work uniquely demonstrates the highly dynamic nature of OTS performance often overlooked during OTS development and deployment. The balance of tRNA in the tRNA pool is particularly integral to maintaining translational fidelity and regulation. Across a wide range of cellular contexts, alteration of the constituency of tRNA isoacceptors has been reported to increase misaminoacylation resulting in protein mistranslation and a decrease in the regulatory capacity of translation^40,67^. Although most tRNA misaminoacylation events are corrected through carefully evolved post-transfer aa-tRNA proofreading mechanisms, orthogonal tRNAs are rarely developed in the context of these editing mechanisms and thus may evade them^68–70^.

While our understanding of the consequences of natural perturbations to the tRNA pool has expanded, little has been done outside of the present work to assess the impact of introducing heavily evolved orthogonal tRNAs specifically on host physiology. We demonstrate here, with innovative proteomics techniques, that orthogonal tRNA is able to interact with native aaRS *in vivo* to reduce translational fidelity in a concentration dependent manner. Traditional methods of analyzing tRNA orthogonality fail to address this dynamic property of tRNA recognition and are therefore more susceptible to reductions in OTS fidelity due to cellular perturbation. Our comprehensive analysis strategy centers on an unbiased survey of orthogonal tRNA misaminoacylation via amino acid misincorporation into a mass spectrometry based reporter. The results of this analysis offer unparalleled insight into authentic and relevant interactions of orthogonal tRNA with the cell and can be directly applied to expand our knowledge of both tRNA and OTS biology.

These considerations, as a whole, illustrate the vast complexity of potential OTS:host interactions and highlight the need for detailed analysis of OTS performance and host cell tolerance. Enriching our understanding of these interactions will ultimately enable more informed OTS design practices which will limit the impact of OTSs on host cells while simultaneously improving OTS performance and robustness across a wide range of experimental settings.

### Establishing permanent OTS installation through systematic characterization

Expression of OTS components episomally from plasmid vectors presents a number of challenges to OTS performance and reliability. Plasmid copy number, for example, can vary substantially according to host strain and growth conditions^71^. This makes even the most carefully designed plasmid relatively unpredictable outside of the conditions in which the OTS was initially implemented. To enhance reliability and ease of use, an OTS could theoretically be permanently installed in the cell. OTSs have been genomically integrated previously, but with few considerations for host environment and interactions with host translational machinery^72^. Here we show how variations in OTS component expression can be benchmarked against native host cellular processes while simultaneously monitoring OTS efficiency. Information on how the orthogonal aaRS and tRNA are positioned in the proteome and cellular substrate pool relative to host counterparts will be critical to install an OTS in the genome with the stability of native translational machinery. Any stress created by overexpression of an OTS component may cause a negative selection pressure resulting in the extrication, or inactivation, of the OTS from the genome. Indeed, we and others have observed transposon inactivation of pSerOTSλ (*unpublished observation*). Data-driven design and integration of OTS components to produce levels of expression comparable to those in native substrate pools paired with improvements to component orthogonality should circumvent extraneous stress resulting from OTS installation. Overall, the ability to successfully install OTSs into the translational complement of host cells will greatly enhance access to these important technologies by improving ease of use and overall reliability, making stable genomic integration an attractive target for the future of OTSs.

## Methods

### Cell growth and general techniques

All DNA oligonucleotides with standard purification and desalting and Sanger DNA sequencing services were obtained through the Keck DNA Sequencing facility and Oligonucleotide Resource, Yale University. Unless otherwise stated, all cultures were grown at 37 °C in LB-Lennox medium (LB, 10 g/L bacto tryptone, 5 g/L sodium chloride, 5 g/L yeast extract). LB agar plates were LB plus 16 g/L bacto agar. Antibiotics were supplemented for selection, where appropriate (Kanamycin, 50 μg/mL and Ampicillin, 100 μg/mL). *E. coli* Top10 cells (Invitrogen, Carlsbad, CA) were used for cloning and plasmid propagation. NEBuilder HiFi Assembly Mix and restriction enzymes were obtained from New England BioLabs. Plasmid DNA preparation was carried out with the QIAprep Spin Miniprep Kit (Qiagen). Pre-cast 4–12% (wt/vol) Bis–Tris gels for SDS–PAGE were purchased from Bio-Rad. Phos-Tag™ reagent for hand-cast protein gels was purchased from Wako.

A complete list of strains and plasmids may be found in Supplemental Table 1. All plasmids were transformed into recipient strains by electroporation. Electrocompetent cells were prepared by inoculating 20 mL of LB with 200 μL of saturated culture and growing at 37 °C until reaching an OD_600_ of 0.4. Cells were harvested by centrifugation at 8,000 RPM for 2 min. at 4 °C. Cells pellets were washed twice with 20 mL ice cold 10% glycerol in deionized water (dH_2_O). Electrocompetent pellets were re-suspended in 100 μL of 10% glycerol in dH2O. 50 ng of plasmid was mixed with 50 μL of re-suspended electrocompetent cells and transferred to 0.1 cm cuvettes, electroporated (BioRad GenePulser™, 1.78 kV, 200 Ω, 25 μF), and then immediately resuspended in 600 μL of LB. Transformed cells were recovered at 37 °C for one hour and 100 μL was subsequently plated on appropriate selective medium.

### Construction of OTS vectors

Detailed construction of OTS plasmids may be found in Supplemental Data Attachment 1. Generally, OTS components were amplified from the progenitor OTS, pSerOTSλ (SepOTSλ, Addgene # 68292). The pSerRS (SepRS9) and EF-pSer (EF-Sep21) were amplified from pSerOTSλ and placed under the control of the glnS* promoter and assembled with the kanamycin selection marker and origin of replication from pSerOTSλ by gibson assembly using NEBuilder HiFi DNA Assembly master mix (NEB). tRNA expression cassettes (1x-, 2x-, 4x-, and 6x-tRNA) were constructed from a 2x-tRNA construct under the control of a proK tRNA promoter and terminator and separated by a valU tRNA linker that was ordered as a DNA fragment from Genewiz. tRNA cassettes were assembled into the previously assembled pSerRS-EF-pSer backbone by gibson assembly creating ColE1-based pSerOTSc OTSs. OTS variants with different origins of replication were created by assembly of Rop amplified from pSerOTSλ into the ColE1-based OTSs and p15a variants were created by replacement of ColE1 with p15a origins by gibson assembly.

### Construction of rEcoli^XpS^

Strain modification using a lambda red based strategy was performed as previously described^73^. Briefly, transformants carrying a lambda red plasmid (pKD46) were grown in 20 mL LB cultures with ampicillin and l-arabinose at 30°C to an OD_600_ of ≈0.6 and then made electrocompetent. 10–100 ng of PCR product targeting serB was transfomred into the prepared cells and recovered in 1 mL of LB for 1 h at 37°C. Following recovery, one-half was spread onto LB-agar plates with Kanamycin for selection. serB gene deletion was verified by PCR and selected colonies were purified non-selectively at 37°C remove pKD46. The KAN deletion cassette was excised from the serB locus using FLP recombinase (pCP20).

### Growth characterization

Stationary phase pre-cultures were obtained by overnight growth with shaking at 37°C in 5 mL of LB supplemented with antibiotic for plasmid maintenance, where appropriate. Stationary phase cultures were diluted to an OD_600_ of 0.01 in 250 μL of LB supplemented with appropriate antibiotic. Growth was monitored on a Biotek Synergy H1 plate reader. OD_600_ was recorded at 10-minute intervals for 16 hours at 37 °C with continuous shaking. All data were measured in triplicate. Growth rate was determined for each replicate and replicates were averaged. Since some strains achieved lower maximum cell densities, slope was calculated based on the linear regression of ln(OD_600_) through 4-6 contiguous time points (40-60 minutes) rather than between two pre-determined OD_600_ values.

### Analytical gel and immunoblotting

100 μM Phos-tag™ acrylamide (Wako) within hand-cast 12% acrylamide gels were used for separation of phosphorylated reporter proteins. SDS-PAGE gels (4–15% acrylamide, Bio-Rad) and Phos-tag™ gels were transferred onto PVDF membranes. All gels were visualized by immunoblot. Anti-His immunoblots were performed using 1:2,500 diluted rabbit Anti-6xHis antibody (PA1-983B, Thermo Fisher Scientific) in 5% w/v milk in TBST for 1 h. Phospho-mTOR (Ser2448) (Cell Signaling, D9C2) Rabbit mAb was used at 1:1,000 dilutions in 5% w/v milk in TBST for 1 h. Phospho-MEK1 production was confirmed with a commercially available phospho-specific antibody for positions 218 and 222 (Phospho-MEK1/2 (Ser217/221), 9154, Cell Signaling Technology) at 1:1,000 dilution in 5% w/v milk in TBST for 1 h. Secondary antibody incubations used 1:10,000 diluted donkey anti-rabbit HRP (711-035-152, Jackson ImmunoResearch) in 5% w/v milk in TBST for 1 h. Protein bands were then visualized using Clarity ECL substrate (Bio-Rad) and an Amersham Imager 600 (GE Healthcare Life Sciences).

### MS-READ reporter purification

Frozen *E. coli* cell pellets were thawed on ice and pellets were lysed by sonication with lysis buffer consisting of 50 mM Tris-HCl (pH 7.4, 23°C), 500 mM NaCl, 0.5 mM EGTA, 1mM DTT, 10 % glycerol, 50 mM NaF, and 1 mM Na_3_O_4_V. The extract was clarified with two rounds of centrifugation performed for 20 minutes at 4 °C and 14,000 x g. Cell free extracts were applied to Ni-NTA metal affinity resin and purified according to the manufacturer’s instructions. Wash buffers contained 50 mM Tris pH 7.5, 500 mM NaCl, 0.5 mM EGTA, 1mM DTT, 50 mM NaF, 1 mM Na_3_VO_4_ and increasing concentrations of imidazole 20 mM, 40mM, and 60mM, sequentially. Proteins were eluted with wash buffer containing 250 mM imidazole. Eluted protein was subjected to 4 rounds of buffer exchange (20mM Tris pH 8.0 and 100mM NaCl) and concentrated using a 30 kDa molecular weight cutoff spin filter (Amicon).

### Protein digestion and mass spectrometry

#### MS-READ analysis

Affinity purified, buffer exchanged protein was digested and analyzed by mass spectrometry as described previously with some modifications. Briefly, the concentration of protein was determined by UV280 spectroscopy and 5 μg ELP-GFP (MS-READ) reporter from *E. coli* was and dissolved in 12.5 μl solubilization buffer consisting of 10 mM Tris-HCl pH=8.5 (23°C), 10 mM DTT, 1 mM EDTA and 0.5 % acid labile surfactant (ALS-101, Protea). Samples were heat denatured for 20 min at 55 °C in a heat block. Alkylation of cysteines was performed with iodoacetamide (IAM) using a final IAM concentration of 24 mM. The alkylation reaction proceeded for 30 min at room temperature in the dark. Excess IAM was quenched with DTT and the buffer concentration was adjusted using a 1 M Tris-HCl pH 8.5 resulting in a final Tris-HCl concentration of 150 mM. The reaction was then diluted with water and 1 M CaCl_2_ solution to obtain a ALS-101 concentration of 0.045 % and 2 mM CaCl_2_ respectively. Finally, sequencing grade porcine trypsin (Promega) was added to obtain an enzyme/protein ratio of 1/5.3 and the digest was incubated for 15 h at 37 °C without shaking. The digest was quenched with 20% TFA solution resulting in a sample pH of 2. Cleavage of the acid cleavable detergent proceeded for 15 min at room temperature. Digests were frozen at −80 °C until further processing. Peptides were desalted on C_18_ UltraMicroSpin columns (The Nest Group Inc.) essentially following the instructions provided by the manufacturer but using 300 μl elution solvent consisting of 80% ACN, 0.1% TFA for peptide elution. Peptides were dried in a vacuum centrifuge at room temperature.

### Dried peptides were reconstituted and analyzed by LC-MS/MS

#### Digestion of intact E. coli for shotgun proteomics

20 mL cultures were inoculated to a starting OD 600nm of 0.01 in LB media using an overnight culture to stationary phase. After reaching mid-log, cells chilled on ice and pelleted by centrifugation for 2 min at 8000 rpm. The resulting pellet was frozen at −80 °C for downstream processing. For cell lysis and protein digest, cell pellets were thawed on ice and 2 ul of cell pellet was transferred to a microcentrifuge tube containing 40 μl of lysis buffer (10 mM Tris-HCl pH 8.6, 10 mM DTT, 1 mM EDTA, and 0.5 % ALS). Cells were lysed by vortex for 30 s and disulfide bonds were reduced by incubating the reaction for 30 min. at 55 °C. The reaction was briefly quenched on ice and 16 μl of a 60 mM IAM solution was added. Alkylation of cysteines proceeded for 30 min in the dark. Excess IAM was quenched with 14 μl of a 25 mM DTT solution and the sample was then diluted with 330 μl of 183 mM Tris-HCl buffer pH 8.0 supplemented with 2 mM CaCl_2_. Proteins were digested overnight using 12 μg sequencing grade trypsin. Following digestion, the reaction was then quenched with 12.5 μl of a 20 % TFA solution, resulting in a sample pH<3. Remaining ALS reagent was cleaved for 15 min at room temperature. The sample (~30 μg protein) was desalted by reverse phase clean-up using C_18_ UltraMicroSpin columns. The desalted peptides were dried at room temperature in a rotary vacuum centrifuge and reconstituted in 30 μl 70 % formic acid 0.1 % TFA (3:8 v/v) for peptide quantitation by UV_280_. The sample was diluted to a final concentration of 0.2 μg/μl and 5 μl (1 μg) was injected for LC-MS/MS analysis.

#### Data acquisition and analysis

LC-MS/MS was performed using an ACQUITY UPLC M-Class (Waters) and Q Exactive Plus mass spectrometer. The analytical column employed was a 65-cm-long, 75-μm-internal-diameter PicoFrit column (New Objective) packed in-house to a length of 50 cm with 1.9 μm ReproSil-Pur 120 Å C18-AQ (Dr. Maisch) using methanol as the packing solvent. Peptide separation was achieved using mixtures of 0.1% formic acid in water (solvent A) and 0.1% formic acid in acetonitrile (solvent B) with either a 90-min gradient 0/1, 2/7, 60/24, 65/48, 70/80, 75/80, 80/1, 90/1; (min/%B, linear ramping between steps). Gradient was performed with a flowrate of 250 nl/min. At least one blank injection (5 μl 2% B) was performed between samples to eliminate peptide carryover on the analytical column. 100 fmol of trypsin-digested BSA or 100 ng trypsin-digested wildtype K-12 MG1655 *E. coli* proteins were run periodically between samples as quality control standards. The mass spectrometer was operated with the following parameters: (MS1) 70,000 resolution, 3e6 AGC target, 300–1,700 m/z scan range; (data dependent-MS2) 17,500 resolution, 1e^6^ AGC target, top 10 mode, 1.6 m/z isolation window, 27 normalized collision energy, 90 s dynamic exclusion, unassigned and +1 charge exclusion. Data was searched using Maxquant version 1.6.10.43 with Deamidation (NQ), Oxidation (M), and Phospho(STY) as variable modifications and Carbamidomethyl (C) as a fixed modification with up to 3 missed cleavages, 5 AA minimum length, and 1% FDR against a modified Uniprot *E. coli* database containing custom MS-READ reporter proteins. MS-READ search results were analyzed using Skyline version 20.1.0.31 and proteome search results were analyzed with Perseus version 1.6.2.2.

