## Supplemental Data for "Principles for Systematic Optimization of an Orthogonal Translation System with Enhanced Biological Tolerance"

### Supplemental Methods and Data

**Supplemental Data Contents** **Page Number**

Supplemental Table 1 ……………………………………………………………………………….. 3

Supplemental Table 2 ……………………………………………………………………………….. 4

Supplemental Table 3 ……………………………………………………………………………….. 5

Supplemental Figure 5 …………………………………………………………………………….. 10

Supplemental Figure 6 …………………………………………………………………………….. 11

Supplemental Figure 7 …………………………………………………………………………….. 11

Supplemental Figure 8 …………………………………………………………………………….. 13

Supplemental Materials and Methods ……………………………………………………………… 14

Supplemental References …………………………………………………………………………… 15

Supplemental Table 1:

**Table S1: E. coli strains and plasmids used in this study**

| Strains | Name | Growth Requirements | Reference |
|---|---|---|---|
| | BL21 (DE3) | No AB required | [1] |
| | BL21 (DE3) ΔserB | No AB required | [2] |
| | BL21 (DE3) B-95 | No AB required | [3] |
| | BL21 (DE3) B-95 ΔserB | No AB required | [4] |
| | C321.ΔA (C321 mutS+, λ-, Δ(ybhB-bioAB)::zeoR, ΔprfA) | Requires d-biotin | [5] |
| | rEcoli$^{XpS}$ (C321 mutS+, λ-, Δ(ybhB-bioAB)::zeoR, ΔprfA, ΔserB) | Requires d-biotin | This Study |
| **Plasmids** | **Name** | **Growth Requirements** | **Reference** |
| E29 | pSerOTSλ | Kanamycin, 37 ˚C | [6] |
| G30 | supD tRNA$^{Ser}$ Suppressor | Kanamycin, 37 ˚C | [6] |
| | *pSerOTSc* | | |
| Q73 | ColE1, proK-1x tRNA, glnS*-pSerRS9-EF-pSer21 | Kanamycin, 37 ˚C | This Study |
| | *pSerOTSc - tRNA Variants* | | |
| P73 | ColE1, proK-2x tRNA | Kanamycin, 37 ˚C | This Study |
| P74 | ColE1, proK-4x tRNA | Kanamycin, 37 ˚C | This Study |
| P75 | ColE1, proK-6x tRNA | Kanamycin, 37 ˚C | This Study |
| | *pSerOTSc - ColE1+ Rop tRNA Variants* | | |
| R20 | ColE1 + Rop, proK-1x tRNA | Kanamycin, 37 ˚C | This Study |
| Q58 | ColE1 + Rop, proK-2x tRNA | Kanamycin, 37 ˚C | This Study |
| Q59 | ColE1 + Rop, proK-4x tRNA | Kanamycin, 37 ˚C | This Study |
| Q60 | ColE1 + Rop, proK-6x tRNA | Kanamycin, 37 ˚C | This Study |
| | *pSerOTSc - p15a tRNA Variants* | | |
| R19 | p15a, proK-1x tRNA | Kanamycin, 37 ˚C | This Study |
| P5 | p15a, proK-2x tRNA | Kanamycin, 37 ˚C | This Study |
| P32 | p15a, proK-4x tRNA | Kanamycin, 37 ˚C | This Study |
| P33 | p15a, proK-6x tRNA | Kanamycin, 37 ˚C | This Study |
| | *pSerOTSc - Modified tRNA* | | |
| V70 | ColE1, G2:C71 1x tRNA | Kanamycin, 37 ˚C | This Study |
| | *OTS Components* | | |
| R4 | pSerOTSλ TRC*-pSerRS only | Kanamycin, 37 ˚C | This Study |
| Q81 | pSerOTSc glnS*-pSerRS only | Kanamycin, 37 ˚C | This Study |
| R9 | pSerOTSλ lpp-5x tRNA only | Kanamycin, 37 ˚C | This Study |
| V60 | ColE1, G2:C71 1x tRNA only | Kanamycin, 37 ˚C | This Study |
| R11 | ColE1, proK-2x tRNA only | Kanamycin, 37 ˚C | This Study |
| R12 | ColE1, proK-4x tRNA only | Kanamycin, 37 ˚C | This Study |
| R3 | ColE1, proK-6x tRNA only | Kanamycin, 37 ˚C | This Study |
| | *Recombinant Reporter Expression* | | |
| N54 | split mCherry + 14-3-3β and PSP3-7, IPTG/Ara Inducible, p15a | Ampicillin, 37 ˚C | [7] |
| C9 | E(17)TAG-GFP, aTc Inducible, RSF1030 | Chloramphenicol, 37 ˚C | [6] |
| PSP2-5 | PSP2-5, human mTOR fragment, aTc Inducible, ColE1 + Rop | Ampicillin, 37 ˚C | [7] |
| U61 | MEK1- S218TAG/S222TAG, aTc Inducible, RSF1030 | Chloramphenicol, 37 ˚C | This Study |
| E52 | TAG-MS-READ mass spectrometry reporter, aTc Inducible, p15a | Ampicillin, 37 ˚C | This Study |

aTc = anhydrotetracycline IPTG= Isopropyl β- d-1-thiogalactopyranoside Ara= Arabinose

Supplemental Table 2:

**Table S2: Summary of growth parameters displayed in Figure 1C-E:**

| Host | Ori. Of Rep | tRNA Copy | Avg. Growth Rate | *S.D.* |
|---|---|---|---|---|
| BL21 | | | 0.057 | 0.003 |
| BL21 | p15a | 1x tRNA | 0.033 | 0.002 |
| BL21 | p15a | 2x tRNA | 0.048 | 0.004 |
| BL21 | p15a | 4x tRNA | 0.041 | 0.001 |
| BL21 | p15a | 6x tRNA | 0.039 | 0.002 |
| BL21 | ColE1 + Rop | 1x tRNA | 0.027 | 0.001 |
| BL21 | ColE1 + Rop | 2x tRNA | 0.022 | 0.001 |
| BL21 | ColE1 + Rop | 4x tRNA | *N.V.* | *N.V.* |
| BL21 | ColE1 + Rop | 6x tRNA | *N.V.* | *N.V.* |
| BL21 | ColE1 | 1x tRNA | 0.025 | 0.001 |
| BL21 | ColE1 | 2x tRNA | 0.030 | 0.001 |
| BL21 | ColE1 | 4x tRNA | *N.V.* | *N.V.* |
| BL21 | ColE1 | 6x tRNA | *N.V.* | *N.V.* |
| **Host** | **Ori. Of Rep** | **tRNA Copy** | **Avg. Growth Rate** | ***S.D.*** |
| rEcoli$^{XpS}$ | | | 0.040 | 0.007 |
| rEcoli$^{XpS}$ | pSerOTSλ | 5x tRNA | 0.015 | 0.002 |
| rEcoli$^{XpS}$ | p15a | 1x tRNA | 0.028 | 0.007 |
| rEcoli$^{XpS}$ | p15a | 2x tRNA | 0.022 | 0.001 |
| rEcoli$^{XpS}$ | p15a | 4x tRNA | 0.019 | 0.002 |
| rEcoli$^{XpS}$ | p15a | 6x tRNA | 0.015 | 0.002 |
| rEcoli$^{XpS}$ | ColE1 + Rop | 1x tRNA | 0.026 | 0.004 |
| rEcoli$^{XpS}$ | ColE1 + Rop | 2x tRNA | 0.026 | 0.004 |
| rEcoli$^{XpS}$ | ColE1 + Rop | 4x tRNA | 0.021 | 0.001 |
| rEcoli$^{XpS}$ | ColE1 + Rop | 6x tRNA | 0.028 | 0.003 |
| rEcoli$^{XpS}$ | ColE1 | 1x tRNA | 0.039 | 0.001 |
| rEcoli$^{XpS}$ | ColE1 | 2x tRNA | 0.022 | 0.001 |
| rEcoli$^{XpS}$ | ColE1 | 4x tRNA | 0.014 | 0.001 |
| rEcoli$^{XpS}$ | ColE1 | 6x tRNA | 0.004 | 0.001 |
| **Host** | **Ori. Of Rep** | **Component** | **Avg. Growth Rate** | ***S.D.*** |
| rEcoli$^{XpS}$ | ColE1 | glnS*-pSerRS | 0.021 | 0.002 |
| rEcoli$^{XpS}$ | ColE1 + Rop | TRC*-pSerRS | 0.012 | 0.001 |
| rEcoli$^{XpS}$ | ColE1 | 2x tRNA Only | 0.025 | 0.001 |
| rEcoli$^{XpS}$ | ColE1 | 4x tRNA Only | 0.020 | 0.003 |
| rEcoli$^{XpS}$ | ColE1 | 6x tRNA Only | 0.018 | 0.005 |
| rEcoli$^{XpS}$ | ColE1 + Rop | λ tRNA Only | 0.020 | 0.002 |

**N.V. = Not Viable**

Supplemental Table 3:

Table S3: Peptides with Gly → pSer substitutions in the E. coli proteome

| Sequence | Length | Protein | Gene | Score | Intensity | Missed Cleavages | MS/MS m/z | Charge | m/z | Retention Time |
|---|---|---|---|---|---|---|---|---|---|---|
| LNEL**G**LQFMQ**G**ARFWHVLDASA**G**K | 24 | P76329 | yedP | 52.541 | 1.33E+08 | 1 | 991.7761841 | 3 | 991.1053 | 93.45 |
| DFKLKG**G**VLP**G**EQEIDTVR | 19 | Q46915 | gudX | 44.616 | 13255000 | 2 | 1193.029419 | 2 | 1192.533 | 101.89 |
| **G**TL**G**QDVIDIRTLGSKGVFTFDP**G**FTSTASCESK | 34 | P0ABH7 | gltA | 42.241 | 6189200 | 2 | 1292.577148 | 3 | 1292.236 | 96.084 |
| MT**G**IVKTFD**G**K | 11 | P0A976 | cspF | 52.482 | 3.22E+08 | 1 | 692.8035889 | 2 | 692.8028 | 71.94 |
| VISEAVAAA**GG**KLQ | 14 | Q46800 | xdhB | 40.994 | 50758000 | 1 | 752.4145508 | 2 | 751.8488 | 79.704 |
| IATLLLP**G**I**G**TIHDLK | 16 | P51020 | mhpE | 48.981 | 36999000 | 0 | 932.0004883 | 2 | 931.9933 | 106.06 |
| T**G**L**G**RRIALILVK | 13 | P39414 | ttdT | 47.261 | 19178000 | 2 | 799.453186 | 2 | 799.4514 | 91.432 |
| **G**ISLQVNAGEHIAIL**G**R | 17 | P23886 | cydC | 68.536 | 29870000 | 0 | 968.9760742 | 2 | 968.4763 | 86.927 |
| IETLCRLT**G**K | 10 | P0A738 | moaC | 121.99 | 2.4E+08 | 1 | 642.8224487 | 2 | 642.8227 | 102.2 |
| LTRPRT**G**NGPR | 11 | P0A8J8 | rhlB | 50.607 | | 2 | 659.8416138 | 2 | 659.8406 | 68.065 |
| LYSMYNSAFLDDLTKAM**G**R | 19 | P75801 | yliF | 40.962 | 33665000 | 1 | 765.0105591 | 3 | 764.3423 | 81.847 |
| NL**G**QENFDAAEK | 12 | P25552 | gppA | 44.753 | 1.72E+08 | 0 | 716.7801514 | 2 | 716.7794 | 64.83 |
| NQVLEKL**G**LNSEEQK | 15 | P18390 | yjjA | 48.741 | | 1 | 913.4230957 | 2 | 913.427 | 65.93 |
| NVELLT**G**FSNR | 11 | P76236 | yeaI | 47.869 | 30157000 | 0 | 672.8151855 | 2 | 672.8134 | 71.466 |
| EPIKNEANN**G**LKNTR | 15 | P23869 | ppiB | 63.624 | 36029000 | 2 | 897.4296875 | 2 | 896.9331 | 84.654 |
| AS**G**IPALPWEDCQ | 13 | P69330 | citD | 46.001 | 41994000 | 0 | 1537.645996 | 1 | 1537.639 | 98.059 |
| M**G**KGKGNVEYWVALIQPGK | 19 | P0ADY7 | rplP | 52.693 | | 2 | 1085.545166 | 2 | 1085.542 | 112.21 |
| QVKTQSCVVA**G**KK | 13 | P39346 | idnD | 45.137 | | 2 | 764.3868408 | 2 | 764.3837 | 68.082 |
| AVAA**G**MNPMDLK | 12 | P0A6F5 | groL | 56.432 | 3005300 | 0 | 664.2930298 | 2 | 664.2929 | 91.465 |
| ELESRQP**G**VR | 10 | P29131 | ftsN | 69.628 | 13653000 | 1 | 633.296814 | 2 | 633.2979 | 83.445 |
| GETFA**G**FKQSR | 11 | P39452 | nrdE | 49.418 | | 1 | 661.3015137 | 2 | 661.3005 | 78.103 |
| IISPMT**G**YVSR | 11 | P27303 | emrA | 70.919 | 23921000 | 0 | 667.817749 | 2 | 667.3147 | 64.414 |
| IV**G**YDEIFGRK | 11 | P30128 | greB | 41.029 | 17380000 | 1 | 695.8461304 | 2 | 695.842 | 87.754 |
| QH**G**LQSMPRLRVMLNEK | 17 | P0AAK1 | hycB | 46.319 | 15848000 | 2 | 711.6933594 | 3 | 711.355 | 106.3 |
| **G**MGESNPVTGNTCDNVK | 17 | P0A910 | ompA | 53.034 | 4607400 | 0 | 945.3768311 | 2 | 945.3739 | 73.828 |
| IDLVQE**G**LR | 9 | P0AF06 | motB | 44.767 | 31797000 | 0 | 569.2811279 | 2 | 569.2812 | 65.859 |
| INGATVDVRL**G**NKFR | 15 | P28248 | dcd | 48.44 | 18301000 | 2 | 878.9468384 | 2 | 878.4431 | 101.71 |
| NTLEIVQE**G**VEAR | 13 | P08201 | nirB | 64.04 | 22226000 | 0 | 776.3782959 | 2 | 776.3744 | 83.343 |
| SFEPL**G**LKK | 9 | P37197 | yhjA | 40.103 | 33542000 | 1 | 556.791687 | 2 | 556.7912 | 77.455 |
| YR**G**VFGQRDVVFMSAK | 16 | P77561 | ydeP | 46.352 | 5.36E+08 | 2 | 986.4676514 | 2 | 985.9633 | 73.381 |
| DEVILPYWRQLID**G**IK | 16 | P0AB89 | purB | 49.35 | 3072700 | 1 | 1027.5271 | 2 | 1027.024 | 114.44 |
| IAENIKF**G**AAQ | 11 | P0AFX9 | rseB | 40.045 | 38493000 | 1 | 629.291626 | 2 | 629.2918 | 73.661 |
| YL**G**RMHQTGR | 10 | P0C0K3 | rdoA | 53.17 | 3868400 | 1 | 657.7987061 | 2 | 657.2947 | 84.611 |

pSer substitution position in annotated as "**G**" in peptide sequence. Peptide in blue featured in Figure 3C.

Supplemental Figure 1:

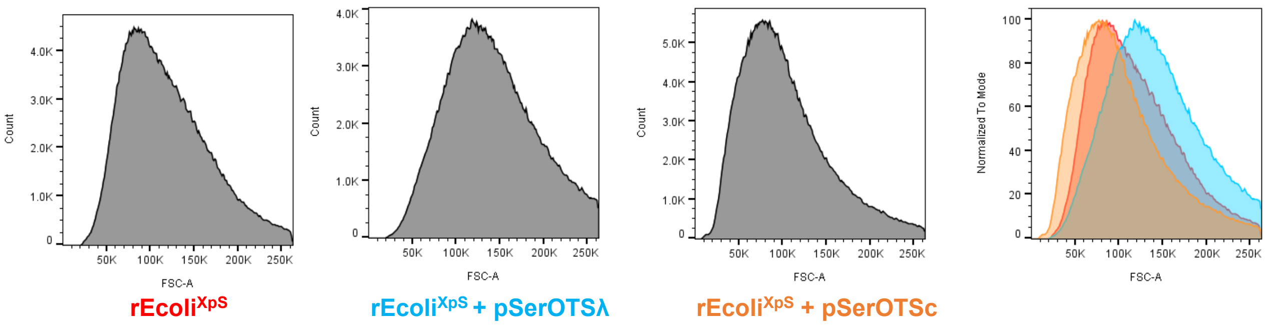

**Figure S1: pSerOTSλ mediates cell size increase in genomically recoded cells.** *E. coli* cell size was measured by flow cytometry with forward light scattering (FSC) used as a proxy for cell size. The size of one million rEcoli$^{XpS}$ cells and cells possessing pSerOTSλ or pSerOTSc were recorded and plotted as histograms representing individual FSC values by abundance in the population. An overlay of sample populations highlights cell size differences across the samples populations; rEcoli$^{XpS}$ (red), rEcoli$^{XpS}$ + pSerOTSc (orange), rEcoli$^{XpS}$ + pSerOTSλ (blue).

Supplemental Figure 2:

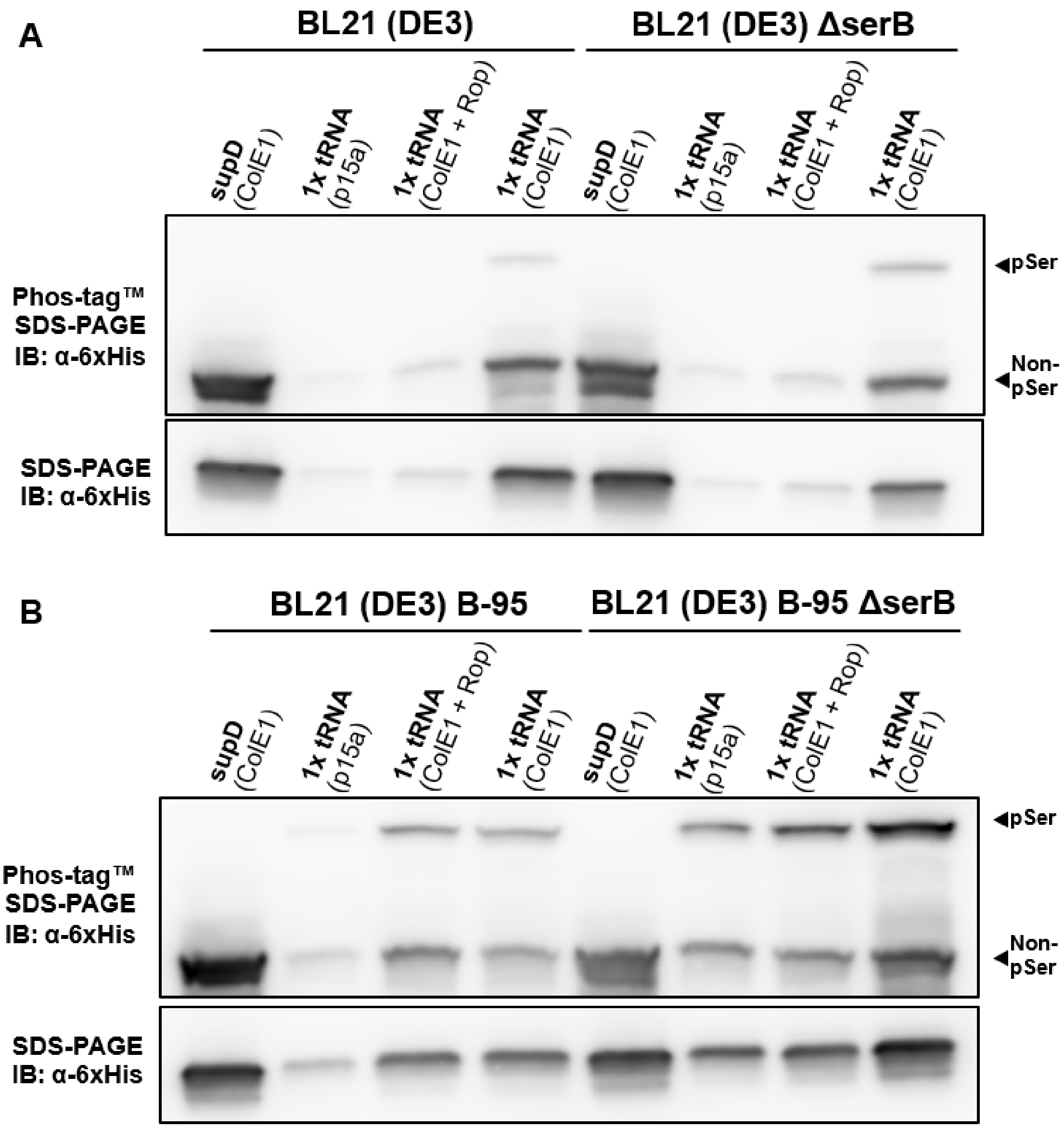

**Figure S2: pSerOTSc performance in BL21 and BL21 B-95.** The performance of 1x tRNA pSerOTSc plasmid copy number variants was assessed by expression of E(17)TAG-GFP reporter in the presence of OTS variants. Crude lysate was separated by SDS-PAGE with and without Phos-tag™ reagent (for phosphoprotein separation) and visualized by immunoblot against 6xHis epitope. Expression and fidelity were assessed in BL21 and BL21 ΔserB (**A**) and in partially recoded BL21 B-95 and BL21 B-95 ΔserB (**B**).

Supplemental Figure 3:

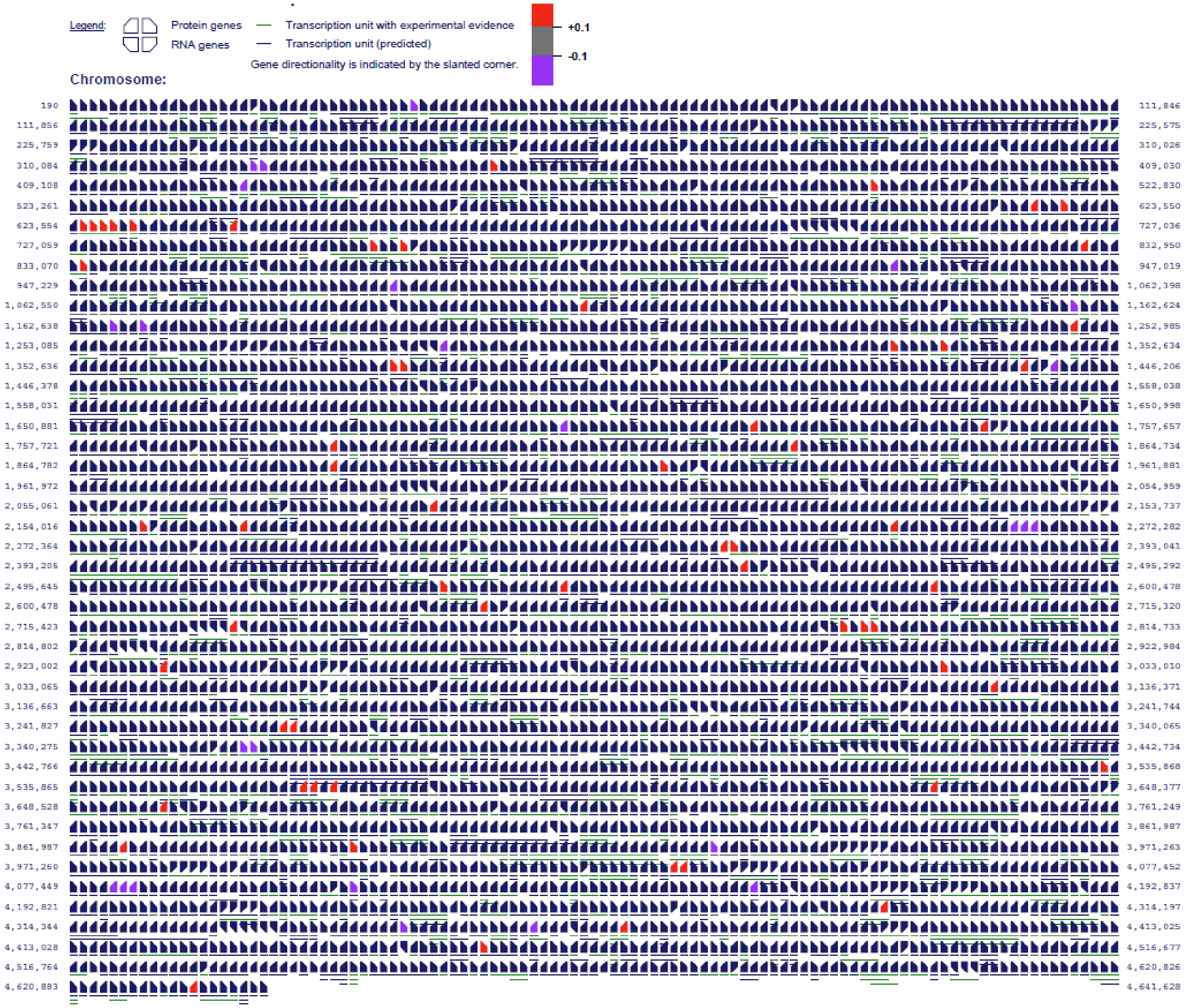

**Figure S3: Alterations to proteome composition mediated by pSerOTSc.** Proteomic analysis of rEcoli$^{XpS}$ cells with and without pSerOTSc was conducted using Perseus. Proteins with statistically significant up-regulation (red) or down-regulation (purple) were matched to their corresponding gene in relation to the complete *E. coli* genome (navy) using Pathway Tools. Proteomes were obtained in triplicate and statistical significance was determined by t-test with p<0.05.

Supplemental Figure 4:

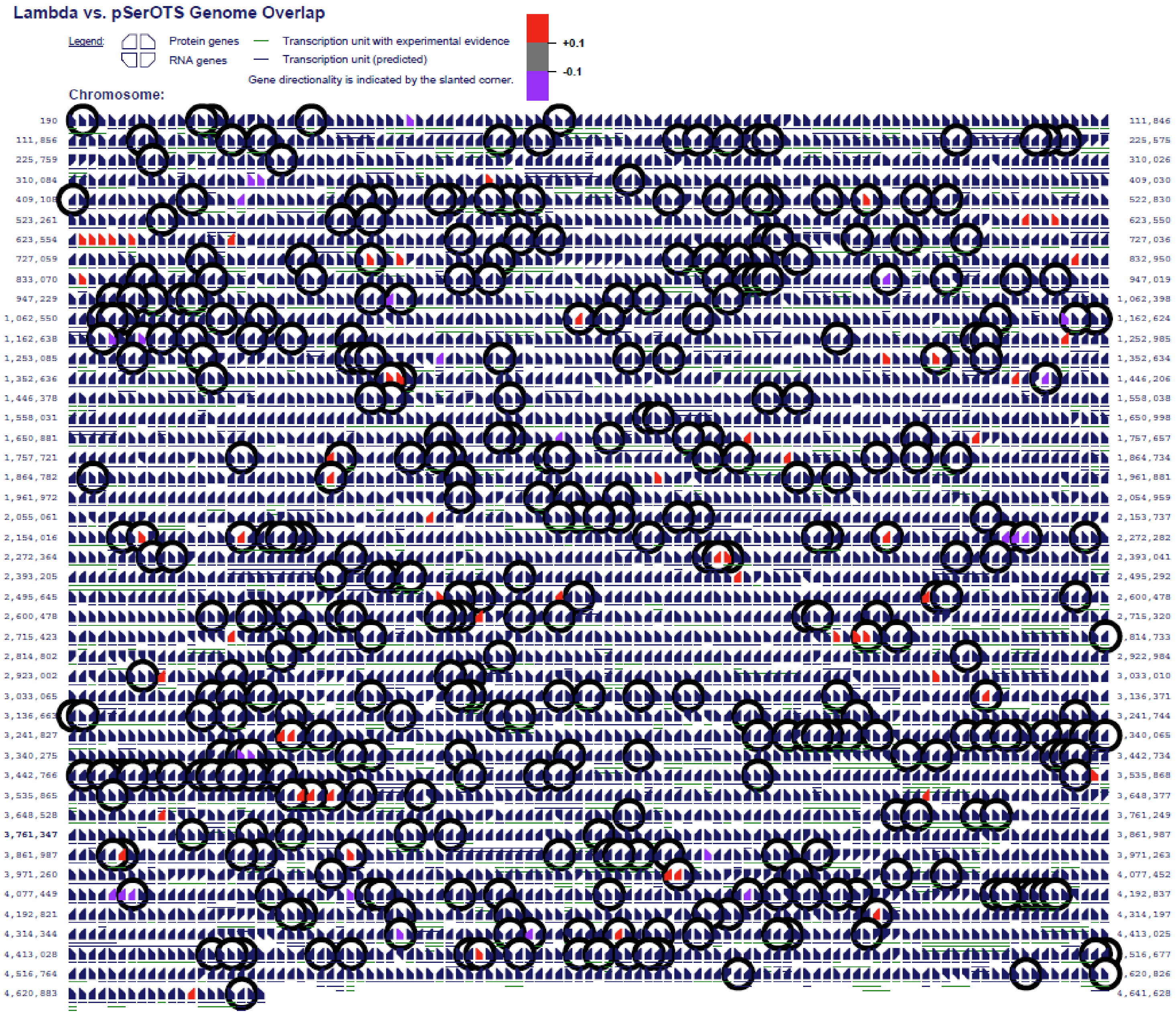

**Figure S4: Alterations to proteome composition mediated by pSerOTSλ.** Proteomic analysis of rEcoli$^{XpS}$ cells with and without pSerOTSλ was conducted using Perseus. Proteins with statistically significant up- or down-regulation in the presence of pSerOTSλ are indicated with black circles, and for pSerOTSc Up-regulation (red) or down-regulation (purple). All proteins were matched to their corresponding gene in relation to the complete *E. coli* genome (navy) using Pathway Tools. Proteomes were obtained in triplicate and statistical significance was determined by t-test with $p<0.05$.

Supplemental Figure 5:

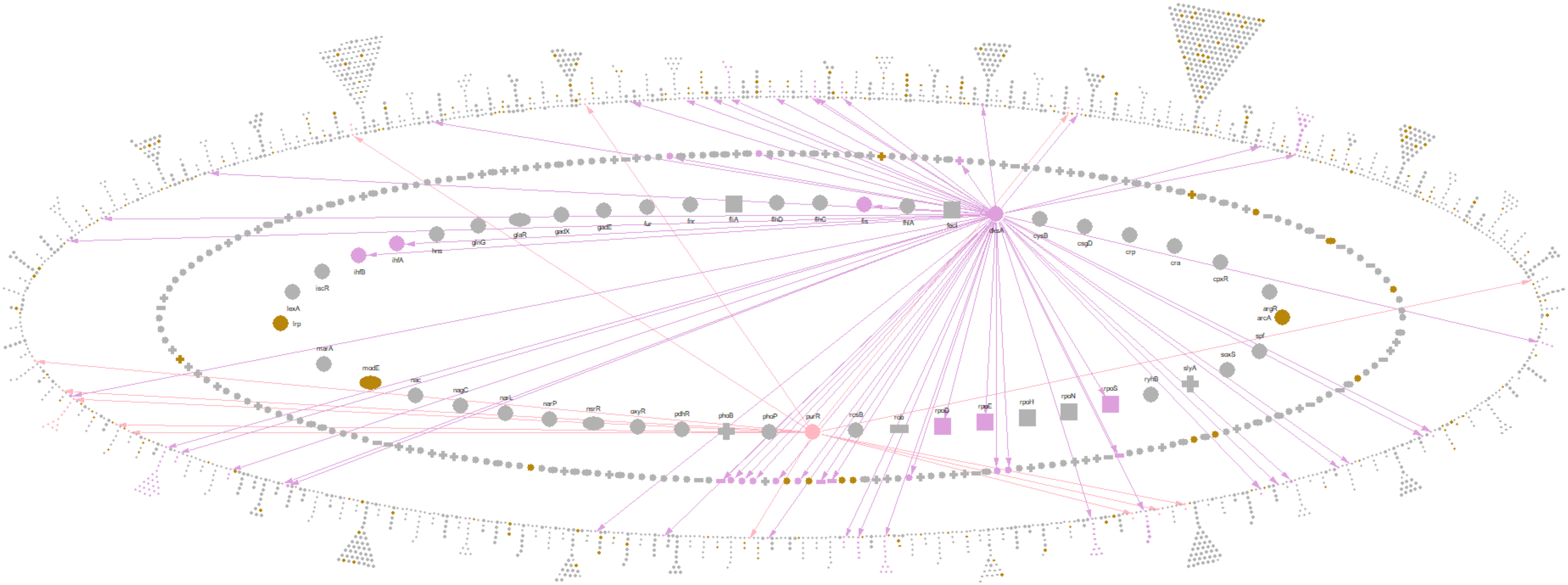

**Figure S5: pSerOTSλ impacts regulatory function in genomically recoded cells.** Alterations to the proteome in cells containing pSerOTSλ were mapped to the E. coli transcriptional regulatory network using Pathway Tools. Two of the most affected regulons dskA and purR, are highlighted in purple and pink, respectively. Arrows colored coded to each node constitute interactions within the regulon. Burnt yellow colored nodes indicate statistically significant change in the protein levels for that specific regulator within the proteome, as compared to WT.

Supplemental Figure 6:

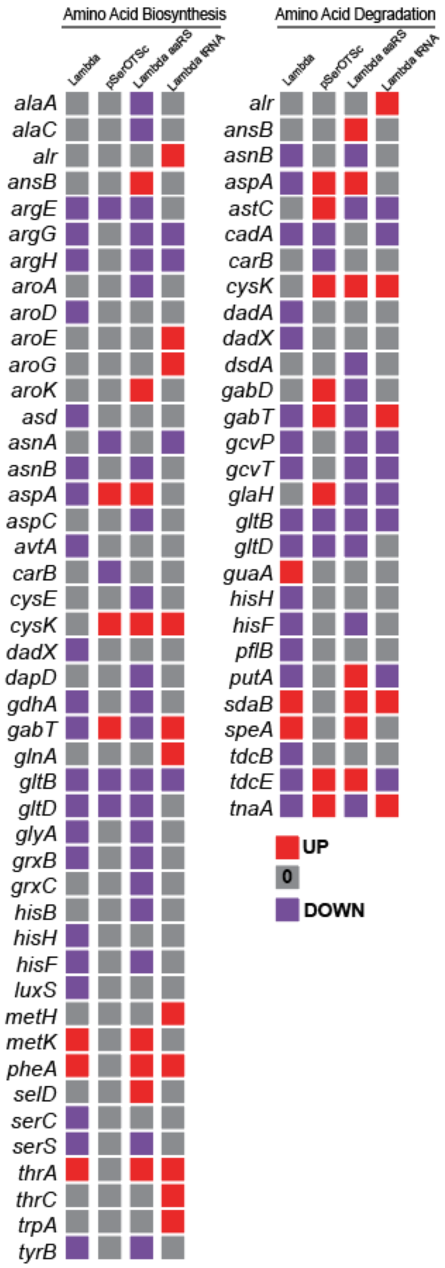

**Figure S6: pSerOTSλ tRNA component expression activates stringent response.** Cells expressing tRNA only were subjected to proteomic analysis and compared to cells expressing full OTSs. Pathway enrichment analysis was conducted for strains harboring OTS variants using Pathway Tools and to illustrate the regulation of the highly impacted pathway components with up-regulated proteins in red, down-regulated proteins in purple, and proteins with no change compared to WT in grey. Enrichment cutoffs were set to a statistically significant differential expression score of 0.1. All proteomes were quantified in triplicate and analyzed in Perseus using t-test and volcano plot functions to obtain statistically significant proteomic deviations.

Supplemental Figure 7:

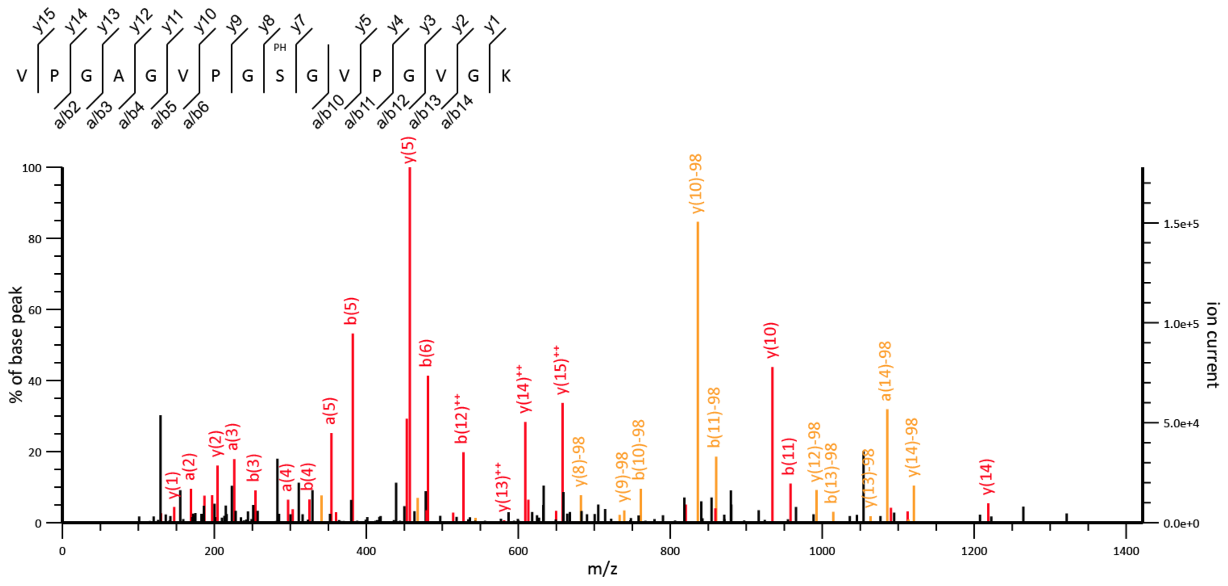

**Figure S7: MS2 ion spectrum and peptide sequence confirming pSer misincorporation.** MS-READ reporter proteins with a central Gly at the guest position were purified from rEcoli$^{XpS}$ cells expressing TRC*-pSerRS alone. Raw mass spectrometry data was processed and searched using Mascot. High confidence identification and peptide sequencing for the Gly-MS-READ reporter peptide (precursor m/z 707.858 M$^{++}$) identified pSer misincorporation at the Gly codon, indicated by PH above Ser between y7 and y8.

Supplemental Figure 8:

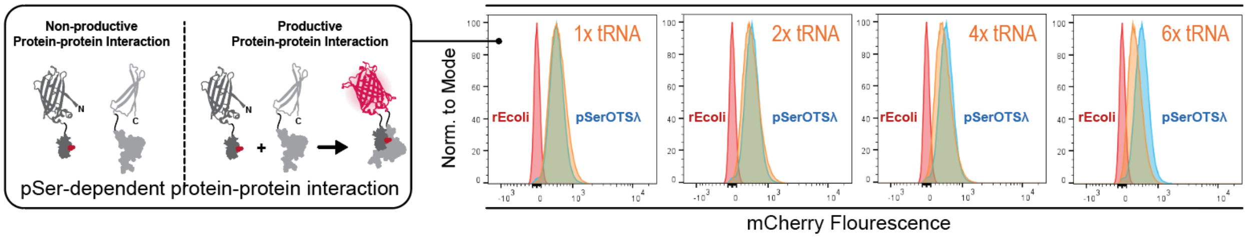

**Figure S8: pSerOTSc variants facilitate phospho-specific protein-protein interactions.** rEcoli$^{XpS}$ cells expressing pSerOTSλ or a pSerOTSc tRNA copy number variant were transformed with a split-mCherry based phospho-specific protein-protein interaction reporter. The C-terminal half of the mCherry reporter was fused to the phospho-binding protein 14-3-3β and the N-terminal half was fused to a phospho-peptide recognized by the phospho-binding protein. Productive interaction mediated by pSer incorporation results in reconstitution of the mCherry flourophore and mCherry fluorescence upon excitation[7]. One million cells were analyzed by flow cytometry and plotted as histograms representing individual mCherry fluorescence values by abundance in the population. When compared to cells containing pSerOTSλ (blue), cells with pSerOTSc displayed comparable fluorescent populations. rEcoli$^{XpS}$ cells (red) without an OTS had no discernable shift in population fluorescence, while pSerOTSc tRNA variants (orange) displayed a tRNA copy number dependent decrease in the magnitude of population fluorescence.

#### Supplemental Methods

*Flow cytometry of cells with HI-P reporter*

5 ml of rEcoli$^{XpS}$ cells containing either the SepOTSλ or pSerOTSc with varying tRNA copy number were diluted to an $OD_{600}$ of 0.15 in 5 ml of LB containing 100 ng/µl ampicillin and 25 ng/µl kanamycin. The cells were grown until $OD_{600}$ reached mid-log (~0.8), then protein expression was induced using 1 mM IPTG, 0.2% arabinose, and 100 ng/µL anhydrotetracycline, where appropriate, and grown at 20 °C and 250 r.p.m. for 20 h. 30-50 µl of cells were mixed with 3 ml ice cold PBS in a 5 ml polystyrene tube (Falcon) for FACS analysis using a BD FACSAria III. For each sample, one million cells were interrogated for mCherry-based fluorescence using a 561-nm laser.
